# Prevalence, phenotype and architecture of developmental disorders caused by *de novo* mutation: The Deciphering Developmental Disorders Study

**DOI:** 10.1101/049056

**Authors:** Jeremy F McRae, Stephen Clayton, Tomas W Fitzgerald, Joanna Kaplanis, Elena Prigmore, Diana Rajan, AlejandroW Sifrim, Stuart Aitken, Nadia Akawi, Mohsan Alvi, Kirsty Ambridge, Daniel M Barrett, Tanya Bayzetinova, Philip Jones, Wendy D Jones, Daniel King, Netravathi Krishnappa, Laura E Mason, Tarjinder Singh, Adrian R Tivey, Munaza Ahmed, Uruj Anjum, Hayley Archer, Ruth Armstrong, Jana Awada, Meena Balasubramanian, Siddharth Banka, Diana Baralle, Angela Barnicoat, Paul Batstone, David Baty, Chris Bennett, Jonathan Berg, Birgitta Bernhard, A Paul Bevan, Maria Bitner-Glindzicz, Edward Blair, Moira Blyth, David Bohanna, Louise Bourdon, David Bourn, Lisa Bradley, Angela Brady, Simon Brent, Carole Brewer, Kate Brunstrom, David J Bunyan, John Burn, Natalie Canham, Bruce Castle, Kate Chandler, Elena Chatzimichali, Deirdre Cilliers, Angus Clarke, Susan Clasper, Jill Clayton-Smith, Virginia Clowes, Andrea Coates, Trevor Cole, Irina Colgiu, Amanda Collins, Morag N Collinson, Fiona Connell, Nicola Cooper, Helen Cox, Lara Cresswell, Gareth Cross, Yanick Crow, Mariella D'Alessandro, Tabib Dabir, Rosemarie Davidson, Sally Davies, Dylan de Vries, John Dean, Charu Deshpande, Gemma Devlin, Abhijit Dixit, Angus Dobbie, Alan Donaldson, Dian Donnai, Deirdre Donnelly, Carina Donnelly, Angela Douglas, Sofia Douzgou, Alexis Duncan, Jacqueline Eason, Sian Ellard, Ian Ellis, Frances Elmslie, Karenza Evans, Sarah Everest, Tina Fendick, Richard Fisher, Frances Flinter, Nicola Foulds, AndrewW Fry, Alan Fryer, Carol Gardiner, Lorraine Gaunt, Neeti Ghali, Richard Gibbons, Harinder Gill, Judith Goodship, David Goudie, Emma Gray, Andrew Green, Philip Greene, Lynn Greenhalgh, Susan Gribble, Lucy Harrison, Victoria Harrison, Rose Hawkins, Liu He, Stephen Hellens, Alex Henderson, Sarah Hewitt, Lucy Hildyard, Emma Hobson, Simon Holden, Muriel Holder, Susan Holder, Georgina Hollingsworth, Tessa Homfray, Mervyn Humphreys, Jane Hurst, Ben Hutton, Stuart Ingram, Melita Irving, Lily Islam, Andrew Jackson, Joanna Jarvis, Lucy Jenkins, Diana Johnson, Elizabeth Jones, Dragana Josifova, Shelagh Joss, Beckie Kaemba, Sandra Kazembe, Rosemary Kelsell, Bronwyn Kerr, Helen Kingston, Usha Kini, Esther Kinning, Gail Kirby, Claire Kirk, Emma Kivuva, Alison Kraus, Dhavendra Kumar, V.K Ajith Kumar, Katherine Lachlan, Wayne Lam, Anne Lampe, Caroline Langman, Melissa Lees, Derek Lim, Cheryl Longman, Gordon Lowther, Sally A Lynch, Alex Magee, Eddy Maher, Alison Male, Sahar Mansour, Karen Marks, Katherine Martin, Una Maye, Emma McCann, Vivienne McConnell, Meriel McEntagart, Ruth McGowan, Kirsten McKay, Shane McKee, Dominic J McMullan, Susan McNerlan, Catherine McWilliam, Sarju Mehta, Kay Metcalfe, Anna Middleton, Zosia Miedzybrodzka, Emma Miles, Shehla Mohammed, Tara Montgomery, David Moore, Sian Morgan, Jenny Morton, Hood Mugalaasi, Victoria Murday, Helen Murphy, Swati Naik, Andrea Nemeth, Louise Nevitt, Ruth Newbury-Ecob, Andrew Norman, Rosie O'Shea, Caroline Ogilvie, Kai-Ren Ong, Soo-Mi Park, Michael J Parker, Chirag Patel, Joan Paterson, Stewart Payne, Daniel Perrett, Julie Phipps, Daniela T Pilz, Martin Pollard, Caroline Pottinger, Joanna Poulton, Norman Pratt, Katrina Prescott, Sue Price, Abigail Pridham, Annie Procter, Hellen Purnell, Oliver Quarrell, Nicola Ragge, Raheleh Rahbari, Josh Randall, Julia Rankin, Lucy Raymond, Debbie Rice, Leema Robert, Eileen Roberts, Jonathan Roberts, Paul Roberts, Gillian Roberts, Alison Ross, Elisabeth Rosser, Anand Saggar, Shalaka Samant, Julian Sampson, Richard Sandford, Ajoy Sarkar, Susann Schweiger, Richard Scott, Ingrid Scurr, Ann Selby, Anneke Seller, Cheryl Sequeira, Nora Shannon, Saba Sharif, Charles Shaw-Smith, Emma Shearing, Debbie Shears, Eamonn Sheridan, Ingrid Simonic, Roldan Singzon, Zara Skitt, Audrey Smith, Kath Smith, Sarah Smithson, Linda Sneddon, Miranda Splitt, Miranda Squires, Fiona Stewart, Helen Stewart, Volker Straub, Mohnish Suri, Vivienne Sutton, Ganesh Jawahar Swaminathan, Elizabeth Sweeney, Kate Tatton-Brown, Cat Taylor, Rohan Taylor, Mark Tein, I Karen Temple, Jenny Thomson, Marc Tischkowitz, Susan Tomkins, Audrey Torokwa, Becky Treacy, Claire Turner, Peter Turnpenny, Carolyn Tysoe, Anthony Vandersteen, Vinod Varghese, Pradeep Vasudevan, Parthiban Vijayarangakannan, Julie Vogt, Emma Wakeling, Sarah Wallwark, Jonathon Waters, Astrid Weber, Diana Wellesley, Margo Whiteford, Sara Widaa, Sarah Wilcox, Emily Wilkinson, Denise Williams, Nicola Williams, Louise Wilson, Geoff Woods, Christopher Wragg, Michael Wright, Laura Yates, Michael Yau, Chris Nellaker, Michael J Parker, Helen V Firth, Caroline F Wright, David R FitzPatrick, Jeffrey C Barrett, Matthew E Hurles

## Abstract

Individuals with severe, undiagnosed developmental disorders (DDs) are enriched for damaging *de novo* mutations (DNMs) in developmentally important genes. We exome sequenced 4,293 families with individuals with DDs, and meta-analysed these data with published data on 3,287 individuals with similar disorders. We show that the most significant factors influencing the diagnostic yield of *de novo* mutations are the sex of the affected individual, the relatedness of their parents and the age of both father and mother. We identified 94 genes enriched for damaging *de novo* mutation at genome-wide significance (P < 7 × 10^−7^), including 14 genes for which compelling data for causation was previously lacking. We have characterised the phenotypic diversity among these genetic disorders. We demonstrate that, at current cost differentials, exome sequencing has much greater power than genome sequencing for novel gene discovery in genetically heterogeneous disorders. We estimate that 42% of our cohort carry pathogenic DNMs (single nucleotide variants and indels) in coding sequences, with approximately half operating by a loss-of-function mechanism, and the remainder resulting in altered-function (e.g. activating, dominant negative). We established that most haploinsufficient developmental disorders have already been identified, but that many altered-function disorders remain to be discovered. Extrapolating from the DDD cohort to the general population, we estimate that developmental disorders caused by DNMs have an average birth prevalence of 1 in 213 to 1 in 448 (0.22-0.47% of live births), depending on parental age.

**Abbreviations:**

PTV: Protein-Truncating Variant
DNM: *De Novo* Mutation
DD: Developmental Disorder
DDD: Deciphering Developmental Disorders study

## Main text

Approximately 2-5% of children are born with major congenital malformations and/or manifest severe neurodevelopmental disorders during childhood ^1,2^. While diverse mechanisms can cause such developmental disorders, including gestational infection and maternal alcohol consumption, damaging genetic variation in developmentally important genes has a major contribution. Several recent studies have identified a substantial causal role for DNMs not present in either parent^3–15^. Despite the identification of many developmental disorders caused by DNMs, it is generally accepted that many more such disorders await discovery^15^, and the overall contribution of DNMs to developmental disorders is not known. Moreover, some pathogenic DNMs completely ablate the function of the encoded protein, whereas others alter the function of the encoded protein^16^; the relative contributions of these two mechanistic classes is also not known.

We recruited 4,293 individuals to the Deciphering Developmental Disorders (DDD) study^15^. Each of these individuals was referred with severe undiagnosed developmental disorders and most were the only affected family member. We systematically phenotyped these individuals and sequenced the exomes of these individuals and their parents. Analyses of 1,133 of these trios were described previously^15,17^. We generated a high sensitivity set of 8,361 candidate DNMs in coding or splicing sequence (mean of 1.95 DNMs per proband), while removing systematic erroneous calls (Supplementary Table 1). 1,624 genes contained two or more DNMs in unrelated individuals.

Twenty-three percent of individuals had likely pathogenic protein-truncating or missense DNMs within the clinically curated set of genes robustly associated with dominant developmental disorders^17^. We investigated factors associated with whether an individual had a likely pathogenic DNM in these curated genes (Figure 1 A, Figure 1 B). We observed that males had a lower chance of carrying a likely pathogenic DNM (*P* = 1.8 × 10^−4^; OR 0.75, 0.65 − 0.87 95% CI), as has also been observed in autism^18^. We also observed increased likelihood of having a pathogenic DNM with the extent of speech delay (*P* = 0.00123), but not other indicators of severity relative to the rest of the cohort. Furthermore, the total genomic extent of autozygosity (due to parental relatedness) was negatively correlated with the likelihood of having a pathogenic DNM (*P* = 1.7 × 10^−7^), for every log_10_ increase in autozygous length, the probability of having a pathogenic DNM dropped by 7.5%, likely due to increasing burden of recessive causation (Figure 1 C). Nonetheless, 6% of individuals with autozygosity equivalent to a first cousin union or greater had a plausibly pathogenic DNM, underscoring the importance of considering *de novo* causation in all families.

**Figure 1.**
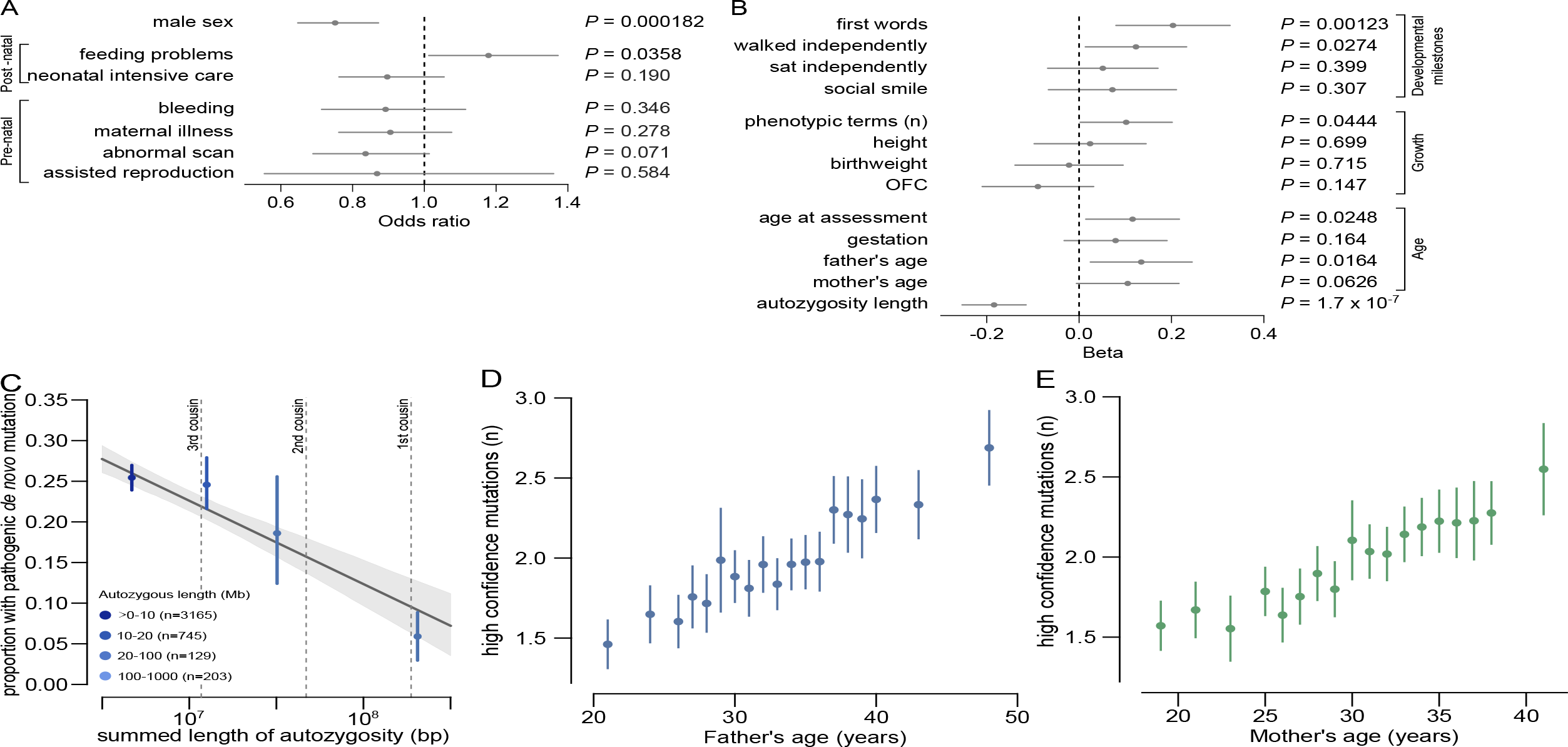
Association of phenotypes with presence of likely pathogenic *de novo* mutations (DNMs). A) Odds ratios and 95% confidence intervals (CI) for binary phenotypes. Positive odds ratios are associated with increased risk of pathogenic DNMs when the phenotype is present. P-values are given for a Fisher's Exact test. B) Beta coefficients and 95% CI from logistic regression of quantitative phenotypes versus presence of a pathogenic DNM. All phenotypes aside from length of autozygous regions were corrected for gender as a covariate. The developmental milestones (age to achieve first words, walk independently, sit independently and social smile) were log-scaled before regression. The growth parameters (height, birthweight and occipitofrontal circumference (OFC)) were evaluated as absolute distance from the median. C) Relationship between length of autozygous regions chance of having a pathogenic DNM. The regression line is plotted as the dark gray line. The 95% confidence interval for the regression is shaded gray. The autozygosity lengths expected under different degrees of consanguineous unions are shown as vertical dashed lines. n, number of individuals in each autozygosity group. D) Relationship between age of fathers at birth of child and number of high confidence DNMs. n, number of high confidence DNMs. E) Relationship between age of mothers at birth of child and number of high confidence DNMs. n, number of high confidence DNMs.

Paternal age has been shown to be the primary factor influencing the number of DNMs in a child^19,20^, and thus is expected to be a risk factor for pathogenic DNMs. Paternal age was only weakly associated with likelihood of having a pathogenic DNM (*P* = 0.016). However, focusing on the minority of DNMs that were truncating and missense variants in known DD-associated genes limits our power to detect such an effect. Analysing all 8,409 high confidence exonic and intronic autosomal DNMs confirmed a strong paternal age effect (*P* = 1.4 × 10^−10^, 1.53 DNMs/year, 1.07−2.01 95% CI), as well as highlighting a weaker, independent, maternal age effect (*P* = 0.0019, 0.86 DNMs/year, 0.32−1.40 95% CI, Figure 1 D, Figure 1 E), as has recently been described in whole genome analyses^21^.

We identified genes significantly enriched for damaging DNMs by comparing the observed gene-wise DNM count to that expected under a null mutation model^22^, as described previously^15^. We combined this analysis with 4,224 published DNMs in 3,287 affected individuals from thirteen exome or genome sequencing studies (Supplementary Table 2)^3–14^ that exhibited a similar excess of DNMs in our curated set of DD-associated genes (Supplementary Figure 1). We found 93 genes with genome-wide significance (*P* < 5 × 10^−7^, Figure 2), 80 of which had prior evidence of DD-association (Supplementary Table 3). We have developed visual summaries of the phenotypes associated with each gene to facilitate clinical use. In addition, we created anonymised average face images from individuals with DNMs in genome-wide significant genes (Figure 2). These images highlight facial dysmorphologies specific to certain genes. To assess any increase in power to detect novel DD-associated genes, we excluded individuals with likely pathogenic variants in known DD-associated genes^15^, leaving 3,158 probands from our cohort, along with 2,955 probands from the meta-analysis studies. In this subset, fourteen genes for which no statistically-compelling prior evidence for DD causation was available achieved genome-wide significance: *CDK13, CHD4, CNOT3, CSNK2A1, GNAI1, KCNQ3, MSL3, PPM1D, PUF60, QRICH1, SET, SUV420H1, TCF20*, and *ZBTB18* (*P* < 5 × 10^−7^, Table 1, Supplementary Figure 4). The clinical features associated with these newly confirmed disorders are summarised in Figure 3, Supplementary Figure 2 and Supplementary Figure 3. *QRICH1* would not achieve genome-wide significance without excluding individuals with likely pathogenic variants in DD-associated genes. In addition to discovering novel DD-associated genes, we identified several new disorders linked to known DD-associated genes, but with different modes of inheritance or molecular mechanisms. We found *USP9X*and *ZC4H2* had a genome-wide significant excess of DNMs in female probands, indicating these genes have X-linked dominant modes of inheritance in addition to previously reported X-linked recessive mode of inheritance in males^23,24^. In addition, we found truncating mutations in *SMC1A* were strongly associated with a novel seizure disorder (*P* = 6.5 × 10^−19^), while in-frame/missense mutations in *SMC1A* with dominant negative effects^25^ are a known cause of Cornelia de Lange Syndrome (CdLS). Individuals with truncating mutations in *SMC1A* lacked the characteristic facial dysmorphology of CdLS.

**Figure 2.**
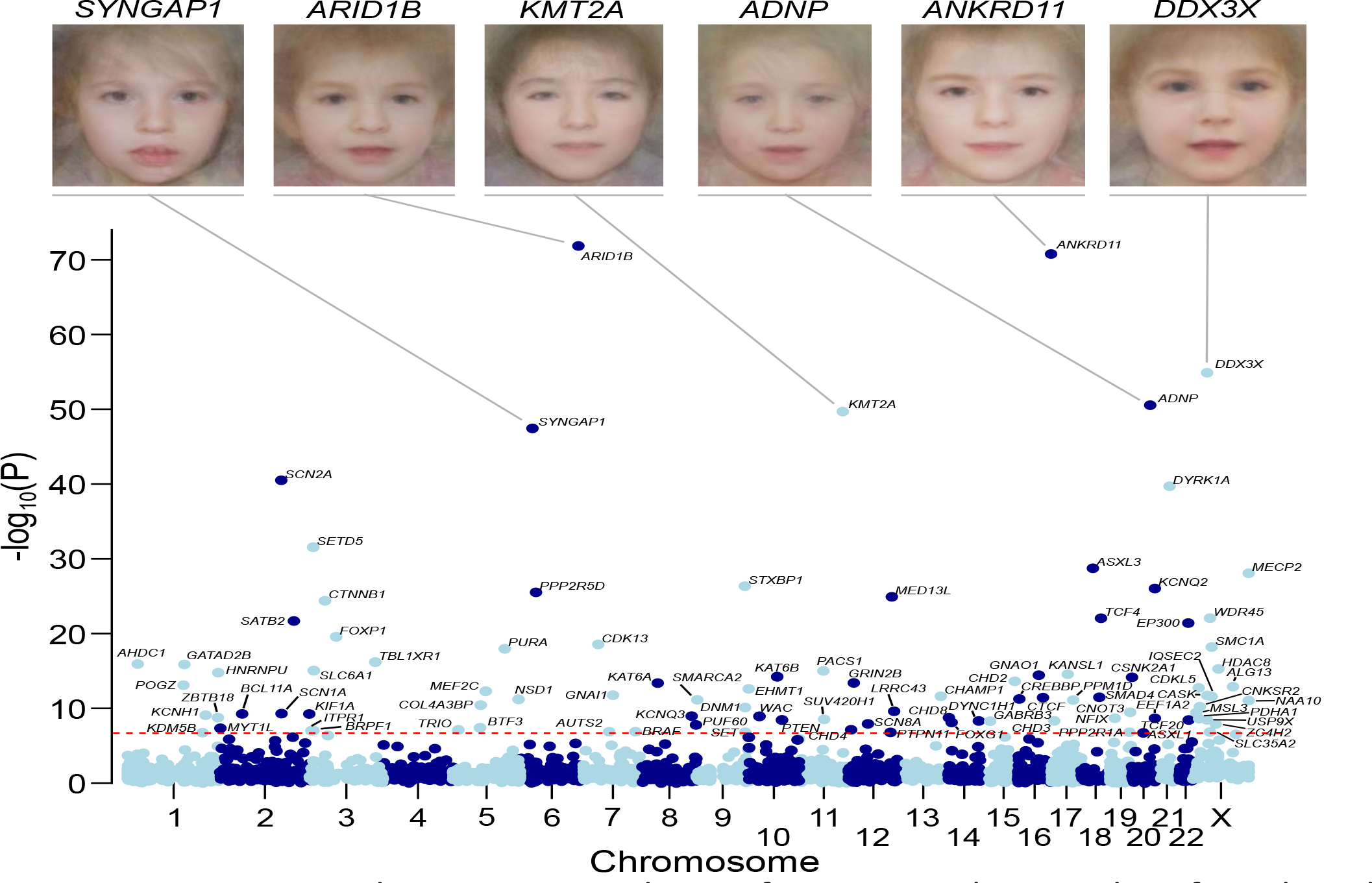
Genes exceeding genome-wide significance. Manhattan plot of combined P-values across all tested genes. The red dashed line indicates the threshold for genome-wide significance (*P* < 7 × 10^−7^). Genes exceeding this threshold have HGNC symbols labelled. Composite facial images from individuals with DNMs in selected genes are included for the six most-significantly associated genes.

**Table 1.** Genes achieving genome-wide significant statistical evidence without previous compelling evidence for being developmental disorder genes. The numbers of unrelated individuals with independent *de novo* mutations (DNMs) are given for protein truncating variants (PTV) and missense variants. If any additional individuals were in other cohorts, that number is given in brackets. The *P*-value reported is the minimum *P*-value from the testing of the DDD dataset or the meta-analysis dataset. The subset providing the *P*-value is also listed. Mutations are considered clustered if the *P*-value proximity clustering of DNMs is less than 0.01.

| Gene | Missense | PTV | P-value | Test | Clustering |
| --- | --- | --- | --- | --- | --- |
| *CDK13* | 10 | 1 | $3.2 \times 10^{-19}$ | DDD | Yes |
| *GNAI1* | 7 (1) | 1 | $2.1 \times 10^{-13}$ | DDD | No |
| *CSNK2A1* | 7 | 0 | $1.4 \times 10^{-12}$ | DDD | Yes |
| *PPM1D* | 0 | 5 (1) | $6.3 \times 10^{-12}$ | Meta | No |
| *CNOT3* | 5 | 2 (1) | $5.2 \times 10^{-11}$ | DDD | Yes |
| *MSL3* | 0 | 4 | $2.2 \times 10^{-10}$ | DDD | No |
| *KCNQ3* | 4 (3) | 0 | $3.4 \times 10^{-10}$ | Meta | Yes |
| *ZBTB18* | 1 (1) | 4 | $1.4 \times 10^{-9}$ | DDD | No |
| *PUF60* | 4 (1) | 3 | $2.6 \times 10^{-9}$ | DDD | No |
| *TCF20* | 1 | 5 | $2.7 \times 10^{-9}$ | DDD | No |
| *SUV420H1* | 0 (2) | 2 (3) | $2.9 \times 10^{-9}$ | Meta | No |
| *CHD4* | 8 (1) | 1 | $7.6 \times 10^{-9}$ | DDD | No |
| *SET* | 0 | 3 | $1.2 \times 10^{-7}$ | DDD | No |
| *QRICH1* | 0 | 3 (1) | $3.6 \times 10^{-7}$ | Meta | No |

**Figure 3.**
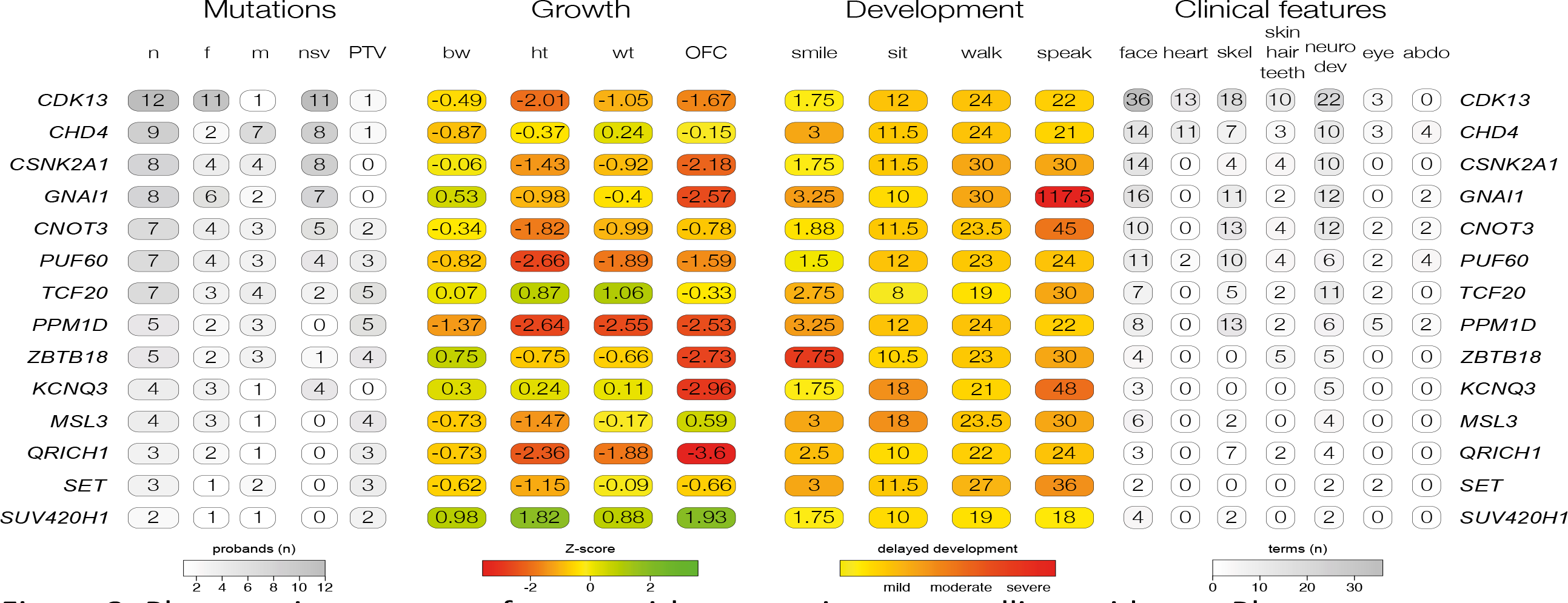
Phenotypic summary of genes without previous compelling evidence. Phenotypes are grouped by type. The first group indicates counts of individuals with DNMs per gene by sex (m: male, f: female), and by functional consequence (nsv: nonsynonymous variant, PTV: protein-truncating variant). The second group indicates mean values for growth parameters: birthweight (bw), height (ht), weight (wt), occipitofrontal circumference (OFC). Values are given as standard deviations from the healthy population mean derived from ALSPAC data. The third group indicates the mean age for achieving developmental milestones: age of first social smile, age of first sitting unassisted, age of first walking unassisted and age of first speaking. Values are given in months. The final group summarises Human Phenotype Ontology (HPO)-coded phenotypes per gene, as counts of HPO-terms within different clinical categories.

We then explored two approaches for integrating phenotypic data into disease gene association: statistical assessment of Human Phenotype Ontology (HPO) term similarity between individuals sharing candidate DNMs in the same gene (as we described previously^26^) and phenotypic stratification based on specific clinical characteristics. Combining genetic evidence and HPO term similarity increased the significance of some known DD-associated genes. However, significance decreased for a larger number of genes causing severe DD but associated with non discriminatory HPO terms (Supplementary Figure 5 A). Although we did not incorporate categorical phenotypic similarity in the gene discovery analyses described above, the systematic acquisition of phenotypic data on affected individuals within DDD enabled aggregate representations to be created for each gene achieving genome-wide significance. We present these in the form of icon-based summaries of growth and developmental milestones (PhenIcons), heatmaps of the recurrently coded HPO terms and, where sufficient face images were available, an anonymised average facial representation (Supplementary Figure 3).

Twenty percent of individuals had HPO terms which indicated seizures and/or epilepsy. We compared analysis within this phenotypically stratified group with gene-wise analyses of the entire cohort, to see if it increased power to detect known seizure-associated genes (Supplementary Figure 5 B). Fifteen seizure-associated genes were genome-wide significant in both the seizure-only and the entire-cohort analyses. Nine seizure-associated genes were genome-wide significant in the entire cohort but not in the seizure subset. Of the 285 individuals with truncating or missense DNMs in known seizure-associated genes, 56% of individuals had no coded terms related to seizures/epilepsy. These findings suggest that the power of increased sample size far outweighs specific phenotypic expressivity due to the shared genetic etiology between individuals with and without epilepsy in our cohort.

The large number of genome-wide significant genes identified in the analyses above allows us to compare empirically different experimental strategies for novel gene discovery in a genetically heterogeneous cohort. We compared the power of exome and genome sequencing to detect genome-wide significant genes, assuming that budget and not samples are limiting, under different scenarios of cost ratios and sensitivity ratios (Figure 4). At current cost ratios (exome costs 30-40% of a genome) and with a plausible sensitivity differential (genome detects 5% more exonic variants than exome^27^) exome sequencing detects more than twice as many genome-wide significant genes. These empirical estimates were consistent with power simulations for identifying dominant loss-of-function genes (Supplementary Figure 6). In summary, while genome sequencing gives greatest sensitivity to detect pathogenic variation in a single individual (or outside of the coding region), exome sequencing is more powerful for novel disease gene discovery (and, analogously, likely delivers lower cost per diagnosis).

**Figure 4.**
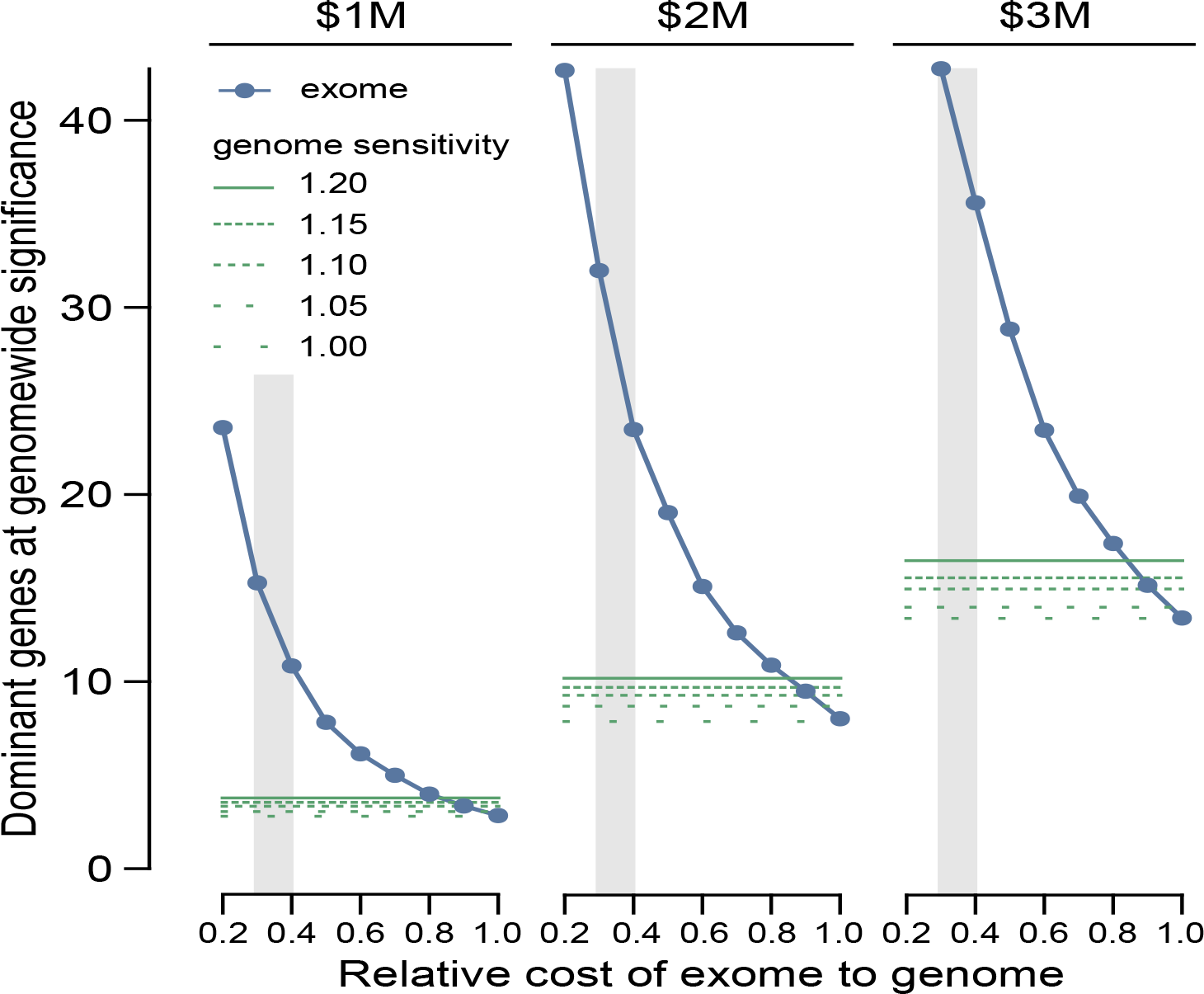
Power of genome versus exome sequencing to discover dominant genes associated with developmental disorders. The power was estimated at three different fixed budgets (1 million (M) USD, 2M and 3M) and a range of relative sensitivities for genomes versus exomes to detect *de novo* mutations. The number of genes identifiable by exome sequencing are shaded blue, whereas the number of genes identifiable by genome sequencing are shaded green. indicate The regions where exome sequencing costs 30-40% of genome sequencing are shaded with a grey background, which corresponds to the price differential in 2016.

Our previous simulations suggested that analysis of a cohort of 4,293 DDD families ought to be able to detect approximately half of all haploinsufficient DD-associated genes at genome-wide significance^15^. Empirically, we have identified 47% (50/107) of haploinsufficient genes previously robustly associated with neurodevelopmental disorders^17^. We hypothesised that genetic testing prior to recruitment into our study may have depleted the cohort of the most clinically recognisable disorders. Indeed, we observed that the genes associated with the most clinically recognisable disorders were associated with a significant, three-fold lower enrichment of truncating DNMs than other DD-associated genes (~40-fold enrichment vs ~120-fold enrichment, Figure 5 A). Removing these most recognisable disorders from the analysis, we identified 55% (42/76) of the remaining haploinsufficient DD-associated genes. The known DD-associated haploinsufficient genes that did not reach genome-wide significance were clearly enriched for those with lower mutability, which we would expect to lower power to detect in our analyses. We identified DD-associated genes (e.g. NRXN2) with high mutability, low clinical recognisability and yet no signal of enrichment for DNMs in our cohort (Supplementary Figure 7). Our analyses call into question whether these genes really are associated with haploinsufficient neurodevelopmental disorders and highlights the potential for well-powered gene discovery analyses to refute prior credence regarding disease gene associations.

**Figure 5.**
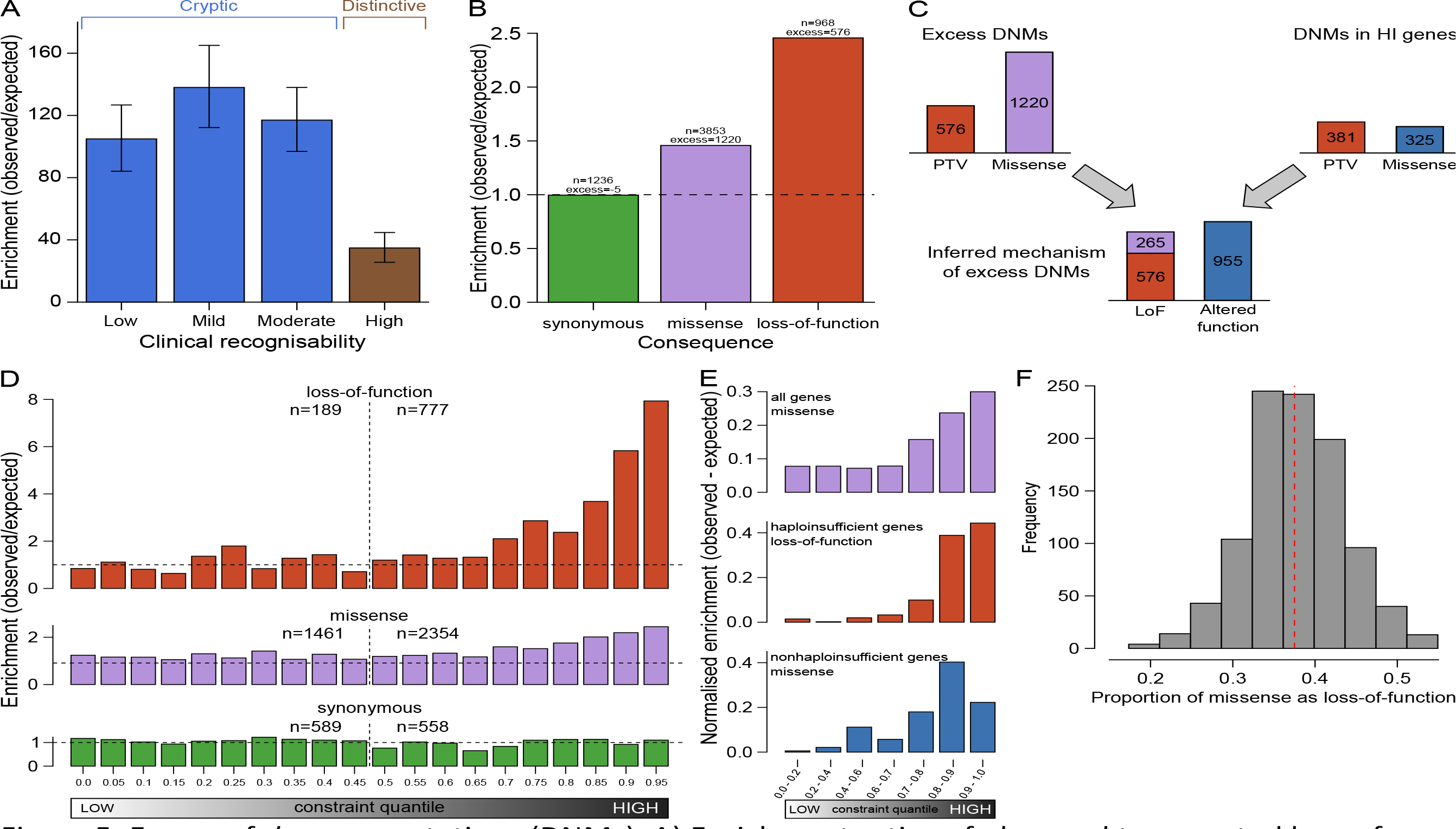
Excess of *de novo* mutations (DNMs). A) Enrichment ratios of observed to expected loss-of-function DNMs by clinical recognisability for dominant haploinsufficient neurodevelopmental genes as judged by two consultant clinical geneticists. B) Enrichment of DNMs by consequence normalised relative to the number of synonymous DNMs. C) Proportion of excess DNMs with loss-of-function or altered-function mechanisms. Proportions are derived from numbers of excess DNMs by consequence, and numbers of excess truncating and missense DNMs in dominant haploinsufficient genes. D) Enrichment ratios of observed to expected DNMs by pLI constraint quantile for loss-of-function, missense and synonymous DNMs. Counts of DNMs in each lower and upper half of the quantiles are provided. E) Normalised excess of observed to expected DNMs by pLI constraint quantile. This includes missense DNMs within all genes, loss-of-function including missense DNMs in dominant haploinsufficient genes and missense DNMs in dominant nonhaploinsufficient genes (genes with dominant negative or activating mechanisms). F) Proportion of excess missense DNMs with a loss-of-function mechanism. The red dashed line indicates the proportion in observed excess DNMs at the optimal goodness-of-fit. The histogram shows the frequencies of estimated proportions from 1000 permutations, assuming the observed proportion is correct.

We estimated the likely prevalence of pathogenic missense and truncating DNMs within our cohort by increasing the stringency of called DNMs until the observed synonymous DNMs equated that expected under the null mutation model (Supplementary Figure 8 A), then quantifying the excess of observed missense and truncating DNMs across all genes (Figure 5 B). We observed an excess of 576 truncating and 1,220 missense mutations, suggesting 41.8% (1,796/4,293) of the cohort has a pathogenic DNM. This estimate of the number of excess missense and truncating DNMs in our cohort is robust to varying the stringency of DNM calling (Supplementary Figure 8 B). The vast majority of synonymous DNMs are likely to be benign, as evidenced by them being distributed uniformly (Figure 5 C) among genes irrespective of their tolerance of truncating variation in the general population (as quantified by the probability of being LoF-intolerant (pLI) metric^28^). By contrast, missense and truncating DNMs are significantly enriched in genes with the highest probabilities of being intolerant of truncating variation (Figure 5 D). Only 51% (923/1,796) of these excess missense and truncating DNMs are located in DD-associated dominant genes, with the remainder likely to affect genes not yet associated with DDs. A much higher proportion of the excess truncating DNMs (71%) than missense DNMs (42%) affected known DD-associated genes. This suggests that whereas most haploinsufficient DD-associated genes have already been identified, many DD-associated genes characterised by pathogenic missense DNMs remain to be discovered.

Understanding the mechanism of action of a monogenic disorder is an important prerequisite for designing therapeutic strategies^29^. We sought to estimate the relative proportion of altered-function and loss-of-function mechanisms among the excess DNMs in our cohort, by assuming that the vast majority of truncating mutations operate by a loss-of-function mechanism and using two independent approaches to estimate the relative contribution of the two mechanisms among the excess missense DNMs (Methods). First, we used the observed ratio of truncating and missense DNMs within haploinsufficient DD-associated genes to estimate the proportion of the excess missense DNMs that likely act by loss-of-function (Figure 5 C). This approach estimated that 47% (42 - 51% 95% CI) of excess missense and truncating DNMs operate by loss-of-function, and 53% by altered-function. Second, we took advantage of the different population genetic characteristics of known altered-function and loss-of-function DD-associated genes. Specifically, we observed that these two classes of DD-associated genes are differentially depleted of truncating variation in individuals without overt developmental disorders (pLI metric^28^). We modelled the observed pLI distribution of excess missense DNMs as a mixture of the pLI distributions of known altered-function and loss-of-function DD-associated genes (Figure 5 E, Figure 5 F), and estimated that 63% (50 - 76% 95% CI) of excess missense DNMs likely act by altered-function mechanisms. Incorporating the truncating DNMs operating by a loss-of-function mechanism, this approach estimated that 57% (48 - 66% 95% CI) of excess missense and truncating DNMs operate by loss-of-function and 43% by altered-function.

We estimated the birth prevalence of monoallelic developmental disorders by using the germline mutation model to calculate the expected cumulative germline mutation rate of truncating DNMs in haploinsufficient DD-associated genes and scaling this upwards based on the composition of excess DNMs in the DDD cohort described above (see Methods), correcting for disorders that are under-represented in our cohort as a result of prior genetic testing (e.g. clinically-recognisable disorders and large pathogenic CNVs identified by prior chromosomal microarray analysis). This gives a mean prevalence estimate of 0.34% (0.31 - 0. 37 95% CI), or 1 in 295 births. By factoring in the paternal and maternal age effects on the mutation rate (Figure 1) we modelled age-specific estimates of birth prevalence (Figure 6) that range from 1 in 448 (both mother and father aged 20) to 1 in 213 (both mother and father aged 45).

**Figure 6.**
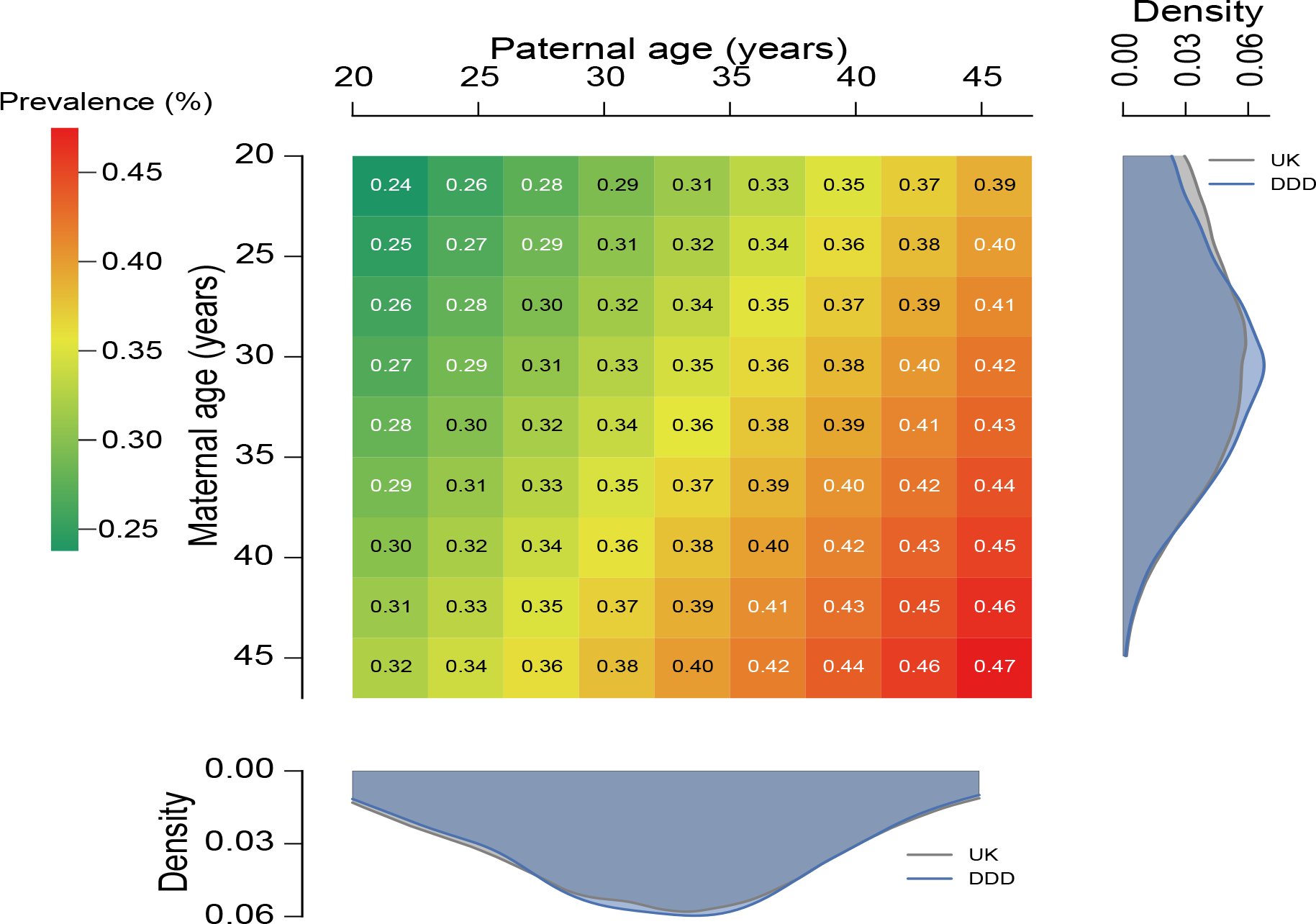
Prevalence of live births with developmental disorders caused by dominant *de novo* mutations (DNMs). The prevalence within the general population is provided as percentage for combinations of parental ages, extrapolated from the maternal and paternal rates of DNMs. Distributions of parental ages within the DDD cohort and the UK population are shown at the matching parental axis.

In summary, we have shown that *de novo* mutations account for approximately half of the genetic architecture of severe developmental disorders, and are split roughly equally between loss-of-function and altered-function. Whereas most haploinsufficient DD-associated genes have already been identified, currently many activating and dominant negative DD-associated genes have eluded discovery. This elusiveness likely results from these disorders being individually rarer, being caused by a relatively small number of missense mutations within each gene. Discovery of the remaining dominant developmental disorders requires larger studies and novel, more powerful, analytical strategies for disease-gene association that leverage gene-specific patterns of population variation, specifically the observed depletion of damaging variation. The integration of accurate and complete quantitative and categorical phenotypic data into the analysis will improve the power to identify ultrarare DD with distinctive clinical presentations. We have estimated the mean birth prevalence of dominant monogenic developmental disorders to be around 1 in 295, which is greater than the combined impact of trisomies 13, 18 and 21^30^ and highlights the cumulative population morbidity and mortality imposed by these individually rare disorders.

## Methods

### Family recruitment

At 24 clinical genetics centers within the United Kingdom (UK) National Health Service and the Republic of Ireland, 4,293 patients with severe, undiagnosed developmental disorders and their parents (4,125 families) were recruited and systematically phenotyped. The study has UK Research Ethics Committee approval (10/H0305/83, granted by the Cambridge South Research Ethics Committee and GEN/284/12, granted by the Republic of Ireland Research Ethics Committee). Families gave informed consent for participation.

Clinical data (growth measurements, family history, developmental milestones, etc.) were collected using a standard restricted-term questionnaire within DECIPHER^31^, and detailed developmental phenotypes for the individuals were entered using Human Phenotype Ontology (HPO) terms^32^. Saliva samples for the whole family and blood-extracted DNA samples for the probands were collected, processed and quality controlled as previously described^15^.

### Exome sequencing

Genomic DNA (approximately 1 μg) was fragmented to an average size of 150 base-pairs (bp) and subjected to DNA library creation using established Illumina paired-end protocols. Adaptor-ligated libraries were amplified and indexed via polymerase chain reaction (PCR). A portion of each library was used to create an equimolar pool comprising eight indexed libraries. Each pool was hybridized to SureSelect ribonucleic acid (RNA) baits (Agilent Human All-Exon V3 Plus with custom ELID C0338371 and Agilent Human All-Exon V5 Plus with custom ELID C0338371) and sequence targets were captured and amplified in accordance with the manufacturer's recommendations. Enriched libraries were subjected to 75-base paired-end sequencing (Illumina HiSeq) following the manufacturer's instructions.

### Alignment and calling single nucleotide variants, insertions and deletions

Mapping of short-read sequences for each sequencing lanelet was carried out using the Burrows-Wheeler Aligner (BWA; version 0.59)^33^ backtrack algorithm with the GRCh37 1000 Genomes Project phase 2 reference (also known as hs37d5). Sample-level BAM improvement was carried out using the Genome Analysis Toolkit (GATK; version 3.1.1)^34^ and SAMtools (version 0.1.19)^35^. This consisted of a realignment of reads around known and discovered indels followed by base quality score recalibration (BQSR), with both steps performed using GATK. Lastly, SAMtools calmd was applied and indexes were created.

Known indels for realignment were taken from the Mills Devine and 1000 Genomes Project Gold set and the 1000 Genomes Project phase low-coverage set, both part of the GATK resource bundle (version 2.2). Known variants for BQSR were taken from dbSNP 137, also part of the GATK resource bundle. Finally, single nucleotide variants (SNVs) and indels were called using the GATK HaplotypeCaller (version 3.2.2); this was run in multisample calling mode using the complete data set. GATK Variant Quality Score Recalibration (VQSR) was then computed on the whole data set and applied to the individual-sample variant calling format (VCF) files. DeNovoGear (version 0.54)^36^ was used to detect SNV, insertion and deletion *de novo* mutations (DNMs) from child and parental exome data (BAM files).

### Variant annotation

Variants in the VCF were annotated with minor allele frequency (MAF) data from a variety of different sources. The MAF annotations used included data from four different populations of the 1000 Genomes Project^37^ (AMR, ASN, AFR and EUR), the UK10K cohort, the NHLBI GO Exome Sequencing Project (ESP), the Non-Finnish European (NFE) subset of the Exome Aggregation Consortium (ExAC) and an internal allele frequency generated using unaffected parents from the cohort.

Variants in the VCF were annotated with Ensembl Variant Effect Predictor (VEP)^38^ based on Ensembl gene build 76. The transcript with the most severe consequence was selected and all associated VEP annotations were based on the predicted effect of the variant on that particular transcript; where multiple transcripts shared the same most severe consequence, the canonical or longest was selected. We included an additional consequence for variants at the last base of an exon before an intron, where the final base is a guanine, since these variants appear to be as damaging as a splice donor variant^26^.

We categorized variants into three classes by VEP consequence:

1. protein-truncating variants (PTV): splice donor, splice acceptor, stop gained, frameshift, initiator codon, and conserved exon terminus variant.
2. missense variants: missense, stop lost, inframe deletion, inframe insertion, coding sequence, and protein altering variant.
3. silent variants: synonymous.

### *De novo* mutation filtering

We filtered candidate DNM calls to reduce the false positive rate but maximize sensitivity, based on prior results from experimental validation by capillary sequencing of candidate DNMs^15^. Candidate DNMs were excluded if not called by GATK in the child, or called in either parent, or if they had a maximum MAF greater than 0.01. Candidate DNMs were excluded when the forward and reverse coverage differed between reference and alternative alleles, defined as *P* < 10^−3^ from a Fisher’s exact test of coverage from orientation by allele summed across the child and parents.

Candidate DNMs were also excluded if they met two of the three following three criteria: 1) an excess of parental alternative alleles within the cohort at the DNMs position, defined as *P* < 10^−3^ under a one-sided binomial test given an expected error rate of 0.002 and the cumulative parental depth; 2) an excess of alternative alleles within the cohort in DNMs in a gene, defined as *P* < 10^−3^ under a one-sided binomial test given an expected error rate of 0.002 and the cumulative depth, or 3) both parents had one or more reads supporting the alternative allele.

If, after filtering, more than one variant was observed in a given gene for a particular trio, only the variant with the highest predicted functional impact was kept (protein truncating > missense > silent). Source code for filtering candidate DNMs can be found here:
https://github.com/jeremymcrae/denovoFilter

### *De novo* mutation validation

For candidate DNMs of interest, primers were designed to amplify 150-250 bp products centered around the site of interest. Default primer3 design settings were used with the following adjustments: GC clamp = 1, human mispriming library used. Site-specific primers were tailed with Illumina adapter sequences. PCR products were generated with JumpStart AccuTaq LA DNA polymerase (Sigma Aldrich), using 40 ng genomic DNA as template. Amplicons were tagged with Illumina PCR primers along with unique barcodes enabling multiplexing of 96 samples. Barcodes were incorporated using Kapa HiFi mastermix (Kapa Biosystems). Samples were pooled and sequenced down one lane of the Illumina MiSeq, using 250 bp paired end reads. An in-house analysis pipeline extracted the read count per site and classified inheritance status per variant using a maximum likelihood approach.

### Individuals with likely pathogenic variants

We previously screened 1,133 individuals for variants that contribute to their disorder^15,17^. All candidate variants in the 1,133 individuals were reviewed by consultant clinical geneticists for relevance to the individuals’ phenotypes. Most diagnosable pathogenic variants occurred *de novo* in dominant genes, but a small proportion also occurred in recessive genes or under other inheritance modes. DNMs within dominant DD-associated genes were very likely to be classified as the pathogenic variant for the individuals’ disorder. Due to the time required to review individuals and their candidate variants, we did not conduct a similar review in the remainder of the 4,293 individuals. Instead we defined likely pathogenic variants as candidate DNMs found in autosomal and X-linked dominant DD-associated genes, or candidate DNMs found in hemizygous DD-associated genes in males. 1,136 individuals in the 4,293 cohort had variants either previously classified as pathogenic^15,17^, or had a likely pathogenic DNM.

### Gene-wise assessment of DNM significance

Gene-specific germline mutation rates for different functional classes were computed^15,22^ for the longest transcript in the union of transcripts overlapping the observed DNMs in that gene. We evaluated the gene-specific enrichment of PTV and missense DNMs by computing its statistical significance under a null hypothesis of the expected number of DNMs given the gene-specific mutation rate and the number of considered chromosomes^22^.

We also assessed clustering of missense DNMs within genes^15^, as expected for DNMs operating by activating or dominant negative mechanisms. We did this by calculating simulated dispersions of the observed number of DNMs within the gene. The probability of simulating a DNM at a specific codon was weighted by the trinucleotide sequence-context^15,22^. This allowed us to estimate the probability of the observed degree of clustering given the null model of random mutations.

Fisher’s method was used to combine the significance testing of missense + PTV DNM enrichment and missense DNM clustering. We defined a gene as significantly enriched for DNMs if the PTV enrichment P-value or the combined missense *P*-value less than 7 × 10^−7^, which represents a Bonferonni corrected *P*-value of 0.05 adjusted for 4x18500 tests (2 × consequence classes tested × protein coding genes).

### Composite face generation

Families were given the option to have photographs of the affected individual(s) uploaded within DECIPHER^31^. Using images of individuals with DNMs in the same gene we generated de-identified realistic average faces (composite faces). Faces were detected using a discriminately trained deformable part model detector^39^. The annotation algorithm identified a set of 36 landmarks per detected face^40^ and was trained on a manually annotated dataset of 3100 images^41^. The average face mesh was created by the Delaunay triangulation of the average constellation of facial landmarks for all patients with a shared genetic disorder.

The averaging algorithm is sensitive to left-right facial asymmetries across multiple patients. For this purpose, we use a template constellation of landmarks based on the average constellations of 2000 healthy individuals^41^. For each patient, we align the constellation of landmarks to the template with respect to the points along the middle of the face and compute the Euclidean distances between each landmark and its corresponding pair on the template. The faces are mirrored such that the half of the face with the greater difference is always on the same side.

The dataset used for this work may contain multiple photos for one patient. To avoid biasing the average face mesh towards these individuals, we computed an average face for each patient and use these personal averages to compute the final average face. Finally, to avoid any image in the composite dominating from variance in illumination between images, we normalised the intensities of pixel values within the face to an average value across all faces in each average. The composite faces were examined manually to confirm successful ablation of any individually identifiable features.

### Assessing power of incorporating phenotypic information

We previously described a method to assess phenotypic similarity by HPO terms among groups of individuals sharing genetic defects in the same gene^26^. We examined whether incorporating this statistical test improved our ability to identify dominant genes at genome-wide significance. Per gene, we tested the phenotypic similarity of individuals with DNMs in the gene. We combined the phenotypic similarity P-value with the genotypic P-value per gene (the minimum P-value from the DDD-only and meta-analysis) using Fisher’s method. We examined the distribution of differences in P-value between tests without the phenotypic similarity P-value and tests that incorporated the phenotypic similarity P-value.

Many (854, 20%) of the DDD cohort experience seizures. We investigated whether testing within the subset of individuals with seizures improved our ability to find associations for seizure specific genes. A list of 102 seizure-associated genes was curated from three sources, a gene panel for Ohtahara syndrome, a currently used clinical gene panel for epilepsy and a panel derived from DD-associated genes^17^. The P-values from the seizure subset were compared to P-values from the complete cohort.

### Assessing power of exome vs genome sequencing

We compared the expected power of exome sequencing versus genome sequencing to identify disease genes. Within the DDD cohort, 55 dominant DD-associated genes achieve genome-wide significance when testing for enrichment of DNMs within genes. We did not incorporate missense DNM clustering due to the large computational requirements for assessing clustering in many replicates.

We assumed a cost of 1,000 USD per individual for genome sequencing. We allowed the cost of exome sequencing to vary relative to genome sequencing, from 10-100%. We calculated the number of trios that could be sequenced under these scenarios. Estimates of the improved power of genome sequencing to detect DNMs in the coding sequence are around 1.05-fold^27^ and we increased the number of trios by 1.0–1.2-fold to allow this.

We sampled as many individuals from our cohort as the number of trios and counted which of the 55 DD-associated genes still achieved genome-wide significance for DNM enrichment. We ran 1000 simulations of each condition and obtained the mean number of genome-wide significant genes for each condition.

### Associations with presence of likely pathogenic *de novo* mutations

We tested whether phenotypes were associated with the likelihood of having a likely pathogenic DNM. Categorical phenotypes (e.g. sex coded as male or female) were tested by Fisher’s exact test while quantitative phenotypes (e.g. duration of gestation coded in weeks) were tested with logistic regression, using sex as a covariate.

We investigated whether having autozygous regions affected the likelihood of having a diagnostic DNM. Autozygous regions were determined from genotypes in every individual, to obtain the total length per individual. We fitted a logistic regression for the total length of autozygous regions on whether individuals had a likely pathogenic DNM. To illustrate the relationship between length of autozygosity and the occurrence of a likely pathogenic DNM, we grouped the individuals by length and plotted the proportion of individuals in each group with a DNM against the median length of the group.

The effects of parental age on the number of DNMs were assessed using 8,409 high confidence (posterior probability of DNM > 0.5) unphased coding and noncoding DNMs in 4,293 individuals. A Poisson multiple regression was fit on the number of DNMs in each individual with both maternal and paternal age at the child’s birth as covariates. The model was fit with the identity link and allowed for overdispersion. This model used exome-based DNMs, and the analysis was scaled to the whole genome by multiplying the coefficients by a factor of 50, based on ~2% of the genome being well covered in our data (exons + introns).

### Excess of *de novo* mutations by consequence

We identified the threshold for posterior probability of DNM at which the number of observed candidate synonymous DNMs equalled the number of expected synonymous DNMs. Candidate DNMs with scores below this threshold were excluded. We also examined the likely sensitivity and specificity of this threshold based on validation results for DNMs within a previous publication^15^ in which comprehensive experimental validation was performed on 1,133 trios that comprise a subset of the families analysed here.

The numbers of expected DNMs per gene were calculated per consequence from expected mutation rates per gene and the 2,407 male and 1,886 females in the cohort. We calculated the excess of DNMs for missense and PTVs as the ratio of numbers of observed DNMs versus expected DNMs, as well as the difference of observed DNMs minus expected DNMs.

### Ascertainment bias within dominant neurodevelopmental genes

We identified 150 autosomal dominant haploinsufficient genes that affected neurodevelopment within our curated developmental disorder gene set. Genes affecting neurodevelopment were identified where the affected organs included the brain, or where HPO phenotypes linked to defects in the gene included either an abnormality of brain morphology (HP:0012443) or cognitive impairment (HP:0100543) term.

The 150 genes were classified for ease of clinical recognition of the syndrome from gene defects by two consultant clinical geneticists. Genes were rated from 1 (least recognisable) to 5 (most recognisable). Categories 1 and 2 contained 5 and 22 genes respectively, and so were combined in later analyses. The remaining categories had more than 33 genes per category. The ratio of observed loss-of-function DNMs to expected loss-of-function DNMs was calculated for each recognisability category, along with 95% confidence intervals from a Poisson distribution given observed counts.

### Proportion of *de novo* mutations with loss-of-function mechanism

The observed excess of missense/inframe indel DNMs is composed of a mixture of DNMs with loss-of-function mechanisms and DNMs with altered-function mechanisms. We found that the excess of PTV DNMs within dominant haploinsufficient DD-associated genes had a greater skew towards genes with high intolerance for loss-of-function variants than the excess of missense DNMs in dominant non-haploinsufficient genes. We binned genes by the probability of being loss-of-function intolerant^28^ constraint decile and calculated the observed excess of missense DNMs in each bin. We modelled this binned distribution as a two-component mixture with the components representing DNMs with a loss-of-function or function-altering mechanism. We identified the optimal mixing proportion for the loss-of-function and altered-function DNMs from the lowest goodness-of-fit (from a spline fitted to the sum-of-squares of the differences per decile) to missense/inframe indels in all genes across a range of mixtures.

The excess of DNMs with a loss-of-function mechanism was calculated as the excess of DNMs with a VEP loss-of-function consequence, plus the proportion of the excess of missense DNMs at the optimal mixing proportion.

We independently estimated the proportions of loss-of-function and altered-function. We counted PTV and missense/inframe indel DNMs within dominant haploinsufficient genes to estimate the proportion of excess DNMs with a loss-of-function mechanism, but which were classified as missense/inframe indel. We estimated the proportion of excess DNMs with a loss-of-function mechanism as the PTV excess plus the PTV excess multiplied by the proportion of loss-of-function classified as missense.

### Prevalence of developmental disorders from dominant *de novo* mutations

We estimated the birth prevalence of monoallelic developmental disorders by using the germline mutation model. We calculated the expected cumulative germline mutation rate of truncating DNMs in 238 haploinsufficient DD-associated genes. We scaled this upwards based on the composition of excess DNMs in the DDD cohort using the ratio of excess DNMs (n=1816) to DNMs within dominant haploinsufficient DD-associated genes (n=412). Around 10% of DDs are caused by *de novo* CNVs^42,43^, which are underrepresented in our cohort as a result of prior genetic testing. If included, the excess DNM in our cohort would increase by 21%, therefore we scaled the prevalence estimate upwards by this factor.

Mothers aged 29.9 and fathers aged 29.5 have children with 77 DNMs per genome on average^20^. We calculated the mean number of DNMs expected under different combinations of parental ages, given our estimates of the extra DNMs per year from older mothers and fathers. We scaled the prevalence to different combinations of parental ages using the ratio of expected mutations at a given age combination to the number expected at the mean cohort parental ages.

## Acknowledgments

We thank the families for their participation and patience. We are grateful to the Exome Aggregation Consortium for making their data available. The DDD study presents independent research commissioned by the Health Innovation Challenge Fund (grant HICF-1009-003), a parallel funding partnership between the Wellcome Trust and the UK Department of Health, and the Wellcome Trust Sanger Institute (grant WT098051). The views expressed in this publication are those of the author(s) and not necessarily those of the Wellcome Trust or the UK Department of Health. The study has UK Research Ethics Committee approval (10/H0305/83, granted by the Cambridge South Research Ethics Committee and GEN/284/12, granted by the Republic of Ireland Research Ethics Committee). The research team acknowledges the support of the National Institutes for Health Research, through the Comprehensive Clinical Research Network. The authors wish to thank the Sanger Human Genome Informatics team, the Sample Management team, the Illumina High-Throughput team, the New Pipeline Group team, the DNA pipelines team and the Core Sequencing team for their support in generating and processing the data. D.R.F. is funded through an MRC Human Genetics Unit program grant to the University of Edinburgh. Finally we gratefully acknowledge the contribution of two esteemed DDD clinical collaborators, John Tolmie and Louise Brueton, who died in the course of the study.

## Author Information

Exome sequencing data are accessible via the European Genome-phenome Archive (EGA) under accession EGAS00001000775. Details of DD-associated genes are available at www.ebi.ac.uk/gene2phenotype. M.E.H. is a co-founder of, and holds shares in, Congenica Ltd, a genetics diagnostic company. Correspondence and requests for materials should be addressed to M.E.H (1015).

## Supplementary Tables

Table provided in external spreadsheet.

Supplementary Table 1: Table of *de novo* mutations (DNM) in the 4,293 DDD individuals. The table includes sex, chromosome, position, reference and alternate alleles, HGNC symbol, VEP consequence, posterior probability of DNM and validation status where available. Individual IDs are available on request. This list excludes the sites that failed validations, but includes sites that passed validation (confirmed), sites that were uncertain (uncertain), and sites that were not tested by secondary validation (NA). Genome positions are given as GRCh37 coordinates.

**Supplementary Table 2:** Details of cohorts used in meta-analyses. This includes numbers of individuals by sex and publication details.

| Phenotype | Year | Male | Female | Note | Citation |
| --- | --- | --- | --- | --- | --- |
| Intellectual disability | 2012 | 47 | 53 |  | De Ligt, et al. <sup>3</sup> |
| Autism spectrum disorder | 2012 | 314 | 29 | subset of lossifov, et al. <sup>9</sup> | lossifov, et al. <sup>10</sup> |
| Autism spectrum disorder | 2012 | 151 | 58 | subset of lossifov, et al. <sup>10</sup> | O’Roak, et al. <sup>11</sup> |
| Intellectual disability | 2012 | 19 | 32 |  | Rauch, et al. <sup>12</sup> |
| Autism spectrum disorder | 2012 | 157 | 68 | subset of lossifov, et al. <sup>9</sup> | Sanders, et al. <sup>13</sup> |
| Seizures | 2013 | 156 | 108 | subset of EuroEPINOMICS-RES Consortium, et al. <sup>6</sup> | Epi4K Consortium and Epilepsy Phenome/Genome Project <sup>5</sup> |
| Congenital heart disease | 2013 | 220 | 142 |  | Zaidi, et al. <sup>14</sup> |
| Seizures | 2014 | 54 | 38 |  | EuroEPINOMICS-RES Consortium, et al. <sup>6</sup> |
| Schizophrenia | 2014 | 308 | 317 |  | Fromer, et al. <sup>7</sup> |
| Intellectual disability | 2014 | 0 | 0 | subset of De Ligt, et al. <sup>3</sup> | Gilissen, et al. <sup>8</sup> |
| Autism spectrum disorder (normal IQ) | 2014 | 1099 | 74 | Counts are for individuals with IQ >= 70. | lossifov, et al. <sup>9</sup> |
| Autism spectrum disorder | 2014 | 446 | 112 | Probands with IQ < 70. | lossifov, et al. <sup>9</sup> |
| Autism spectrum disorder | 2014 | 1192 | 253 | Counts are extrapolated from the sex ratio of individuals with *de novo* mutations. | De Rubeis, et al. <sup>4</sup> |

Supplementary Table 3: Genes with genome-wide significant statistical evidence to be developmental disorder genes. The numbers of unrelated individuals with independent *de novo* mutations (DNMs) are given for protein truncating variants (PTV) and missense variants. If any additional individuals were in other cohorts, that number is given in brackets. The *P*-value reported is the minimum *P*-value from the testing of the DDD dataset or the meta-analysis dataset. The subset providing the *P*-value is also listed. Mutations are considered clustered if the *P*-value proximity clustering of DNMs is less than 0.01.

### Supplementary Figures

**Supplementary Figure 1:**
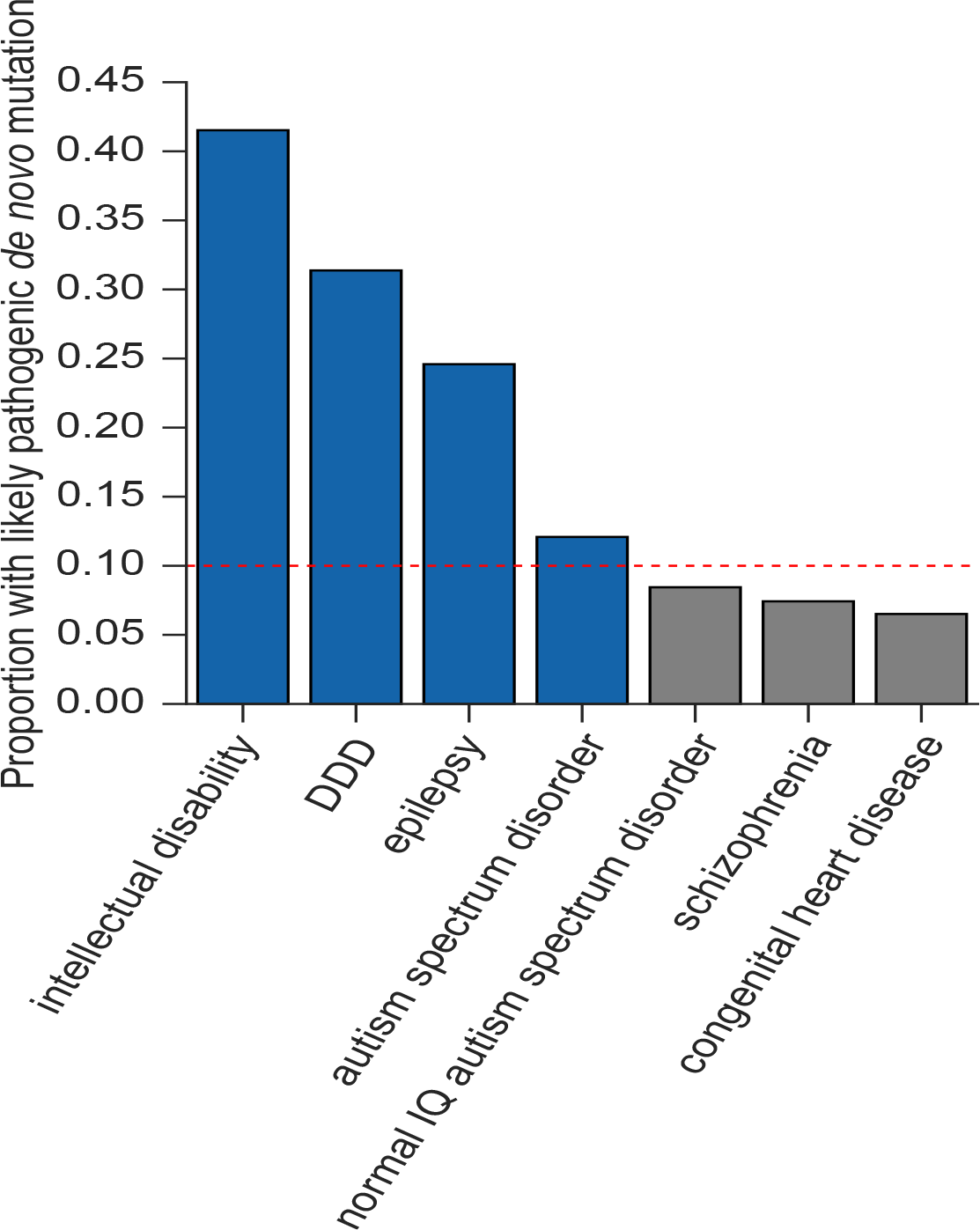
Proportion of individuals with a *de novo* mutation (DNM) likely to be pathogenic. These only included individuals with protein altering or protein truncating DNMs in dominant or X-linked dominant developmental disorder (DD) associated genes, or males with DNMs in hemizygous DD-associated genes. The proportions given are for those individuals with any DNMs rather than the total number of individuals in each subset. Cohorts included in the DNM meta-analyses are shaded blue.

**Supplementary Figure 2:**
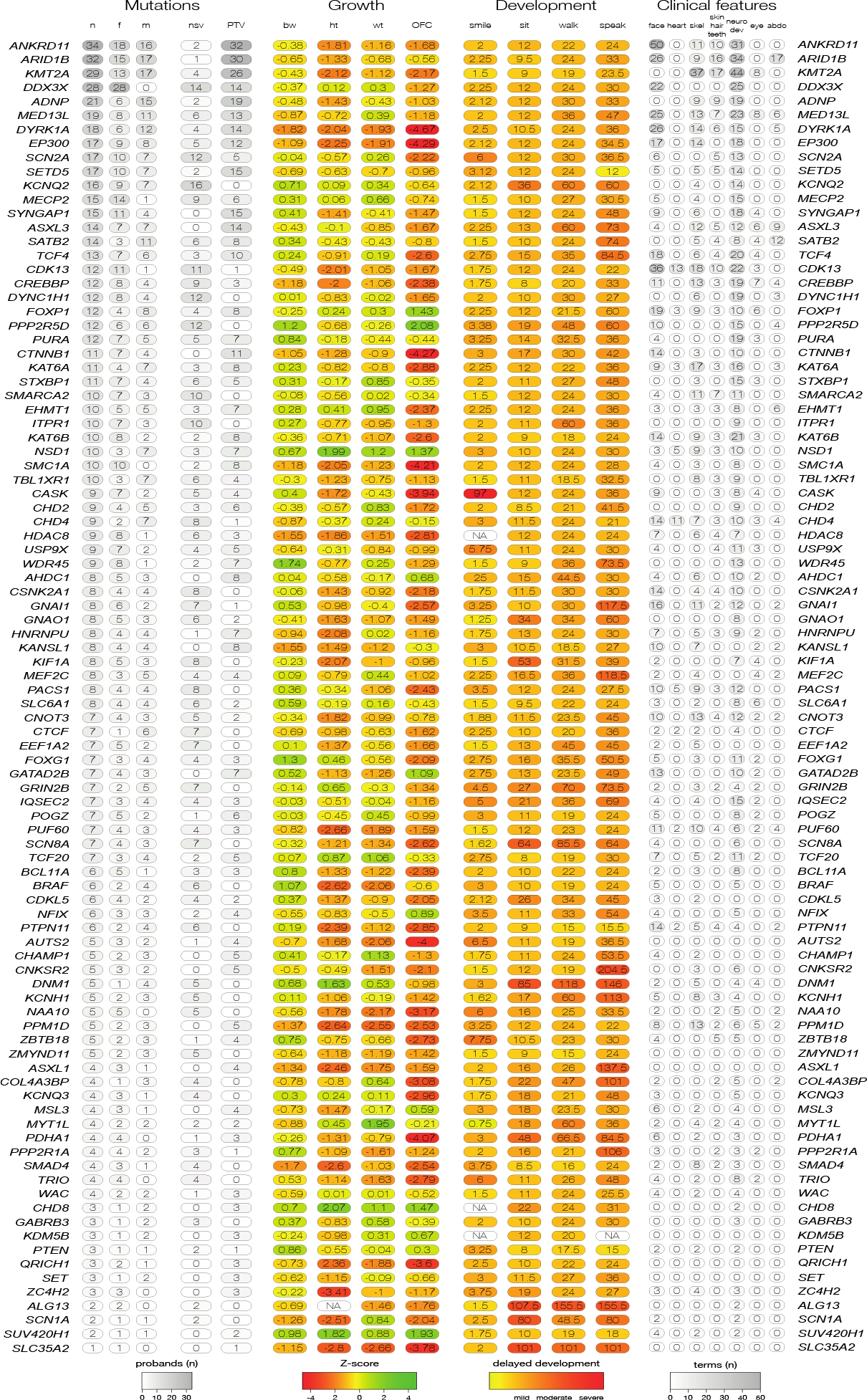
Phenotypic summary of individuals with *de novo* mutations in genes achieving genomewide significance. Phenotypes are grouped by type. The first group indicates counts of individuals with DNMs per gene by sex (m: male, f: female), and by functional consequence (nsv: nonsynonymous variant, PTV: protein-truncating variant). The second group indicates mean values for growth parameters: birthweight (bw), height (ht), weight (wt), occipitofrontal circumference (OFC). Values are given as standard deviations from the healthy population mean derived from ALSPAC data. The third group indicates the mean age for achieving developmental milestones: age of first social smile, age of first sitting unassisted, age of first walking unassisted and age of first speaking. Values are given in months. The final group summarises Human Phenotype Ontology (HPO)-coded phenotypes per gene, as counts of HPO-terms within different clinical categories.

**Supplementary Figure 3:**
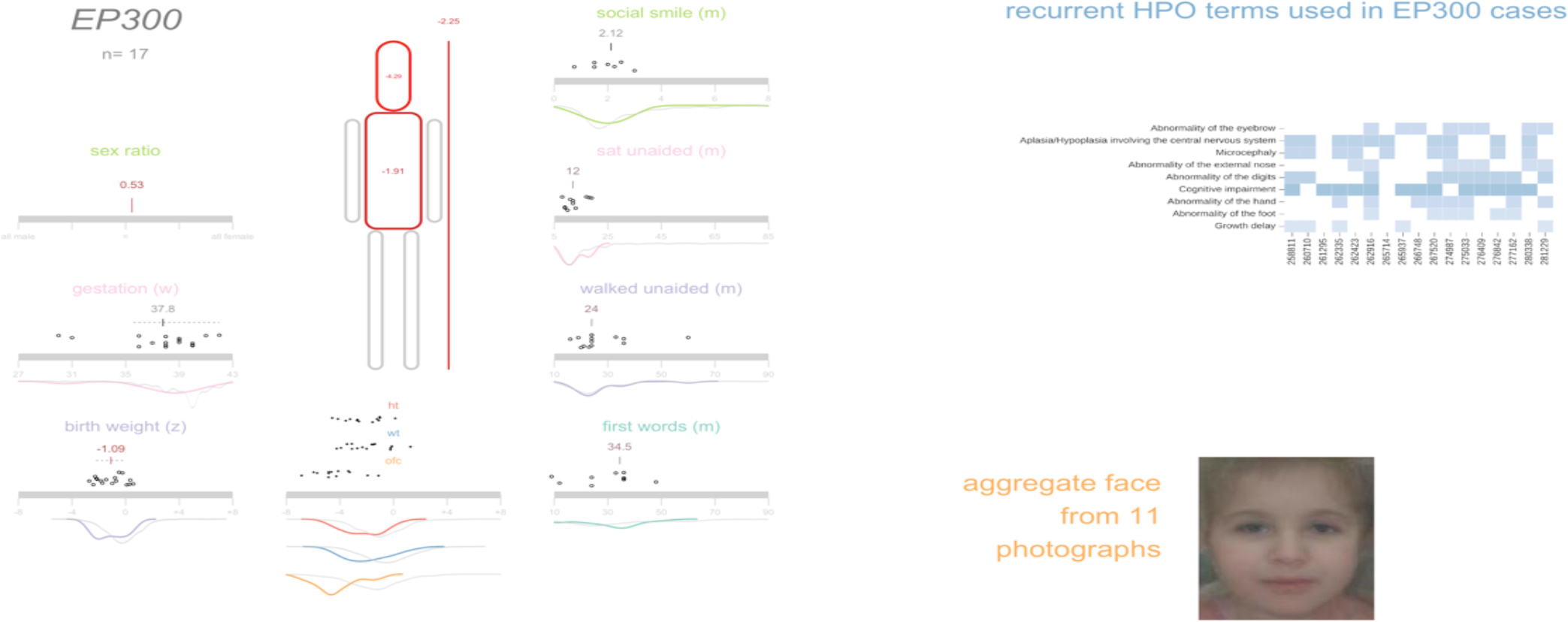
Example of an icon-, heat map-and image-based summary of the quantitative, categorical and average face for each of the genes exceeding genome-wide significance. This uses data on the 17 individuals with de novo mutations (DNMs) in *EP300*. A separate pdf file containing these “phenicons” for all genes is provided. Each has up to three parts. The left hand half of each page provides visual representations of the gene name, the number of individuals with de novo mutations in that gene, sex ratio, gestation (in weeks), anthropometric data (z scores for birth weight, height, weight and occipital-frontal head circumference (ofc)) and developmental milestones (in months for attainment of social smile, sitting unaided, walking unaided and first clear words) from individuals with DNMs in the gene. The scaled cartoon figure shows the height weight and OFC with the colour of the head, trunk and height graded with grey representing a z score of 0 and red increasing negative and green increasing positive scores. For each metric a scatter plot is given above the indicator bar representing the measurement for each individual. Where more that four values are available two density plots are given below the bar the grey representing the data for all individuals in the 94-gene set and coloured the density plot for the gene in question. In *EP300* the OFC measurements are shifted significantly to the left compared to the whole group. For the z score data mean values are provided and for the developmental data median values are given above the bar. The top panel on the left hand side of the page summarises the key Human Phenotype Ontology (HPO) terms for each gene. The HPO terms in the individuals were selected, including the ancestral terms. Terms that are rarer in the 4,293 individuals rank higher, adjusted by the number of individuals with DNMs who had the term. The heatmaps are shaded by the number of individuals with each term. The heatmaps exclude terms that rank lower than a descendant term (excluding more general terms if a more specific term occurred first), and terms where fewer than 25% of individuals had the term, or in genes with less than 8 individuals, terms with fewer than two individuals. The bottom panel on the right hand half of the page summarises the facial photographs from individuals with DNMs in each gene. The averaged face images are only available for selected genes, based on the availability of sufficient high-quality facial photographs of individuals for each gene. The whole image was generated using a custom R script employing grid based graphics.

**Supplementary Figure 4:**
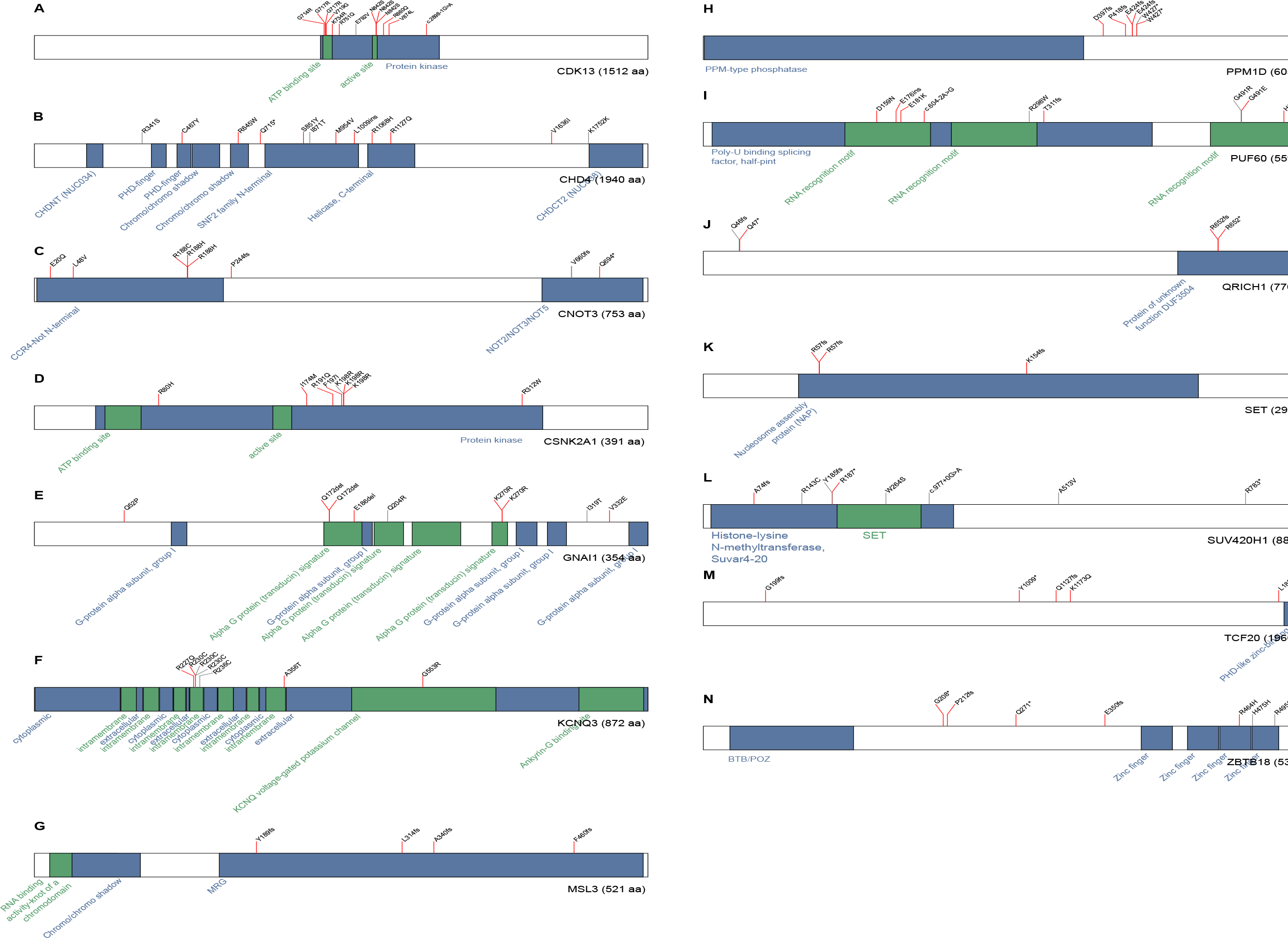
Dispersion of *de novo* mutations and domains for each novel gene. A) *CDK13*, B) *CHD4*, C) *CNOT3*, D) *CSNK2A1,*, E) *GNAI1*, F) *KCNQ3*, G) *MSL3*, H) *PPM1D*, I) *PUF60*, J) *QRICH1*, K) *SET*, L) *SUV420H1*, M) *TCF20* and N) *ZBTB18*.

**Supplementary Figure 5:**
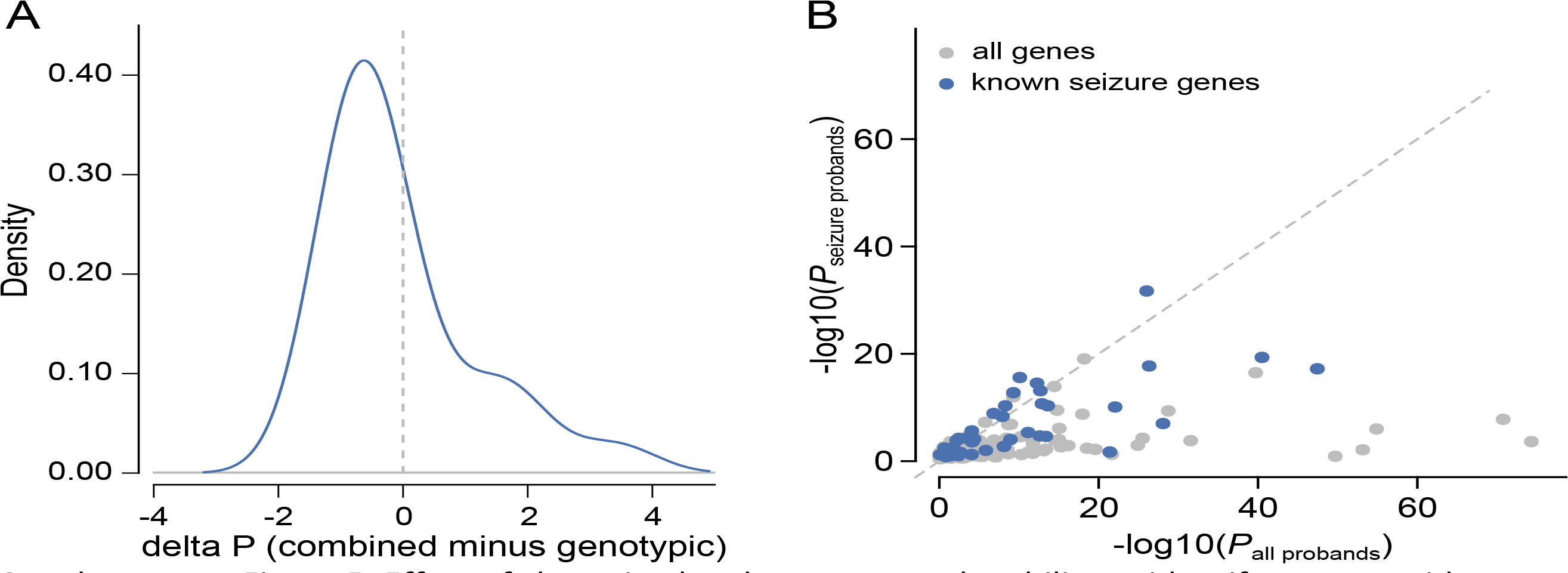
Effect of clustering by phenotype on the ability to identify genomewide significant genes. A) Comparison of P-values derived from genotypic information alone versus *P*-values that incorporate genotypic information and phenotypic similarity. B) Comparison of *P*-values from tests in the complete DDD cohort versus tests in the subset with seizures. Genes that were previously linked to seizures are shaded blue.

**Supplementary Figure 6:**
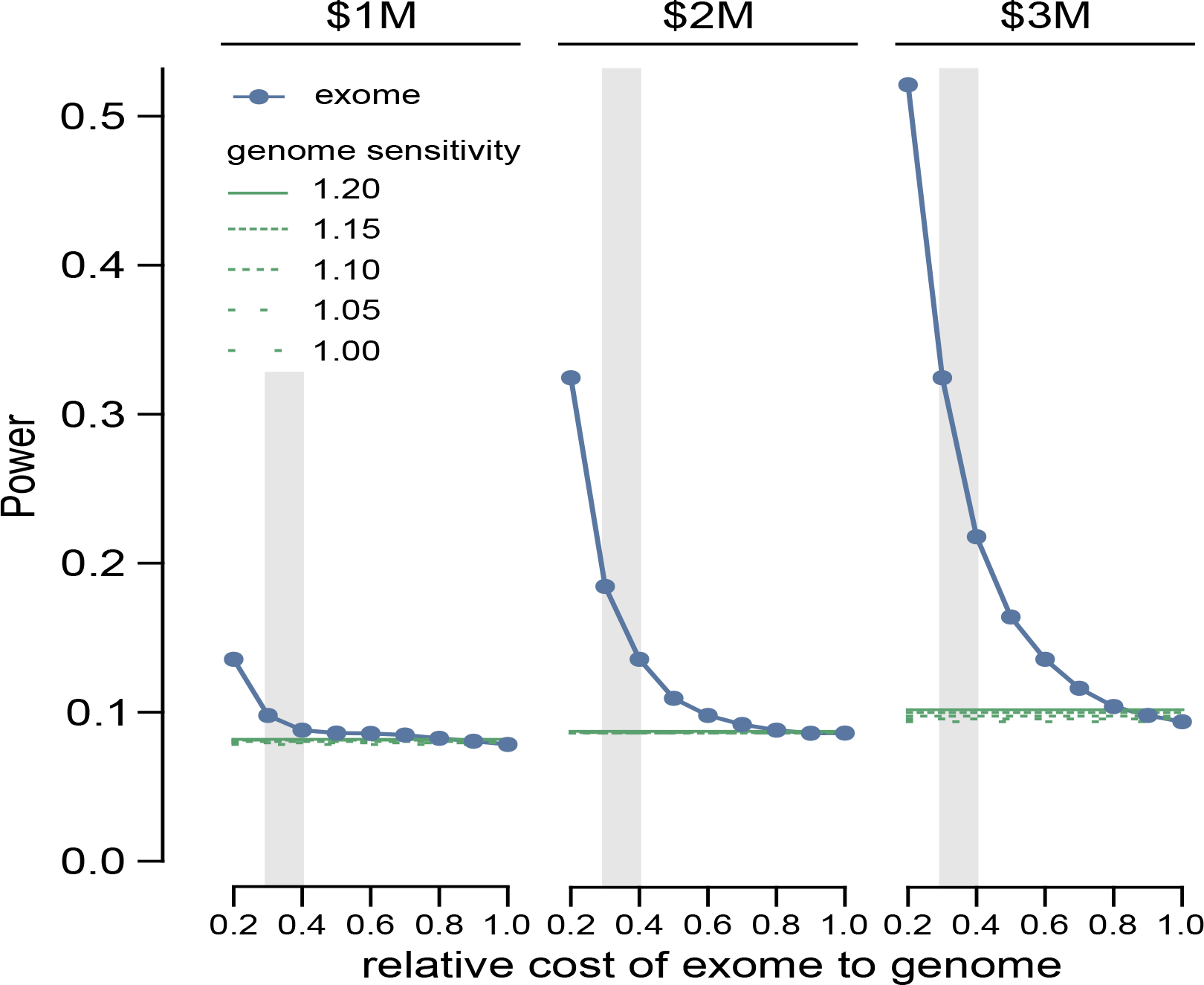
Simulated estimates of power to detect loss-of-function genes in the genome at difference cohort sizes, given fixed budgets.

**Supplementary Figure 7:**
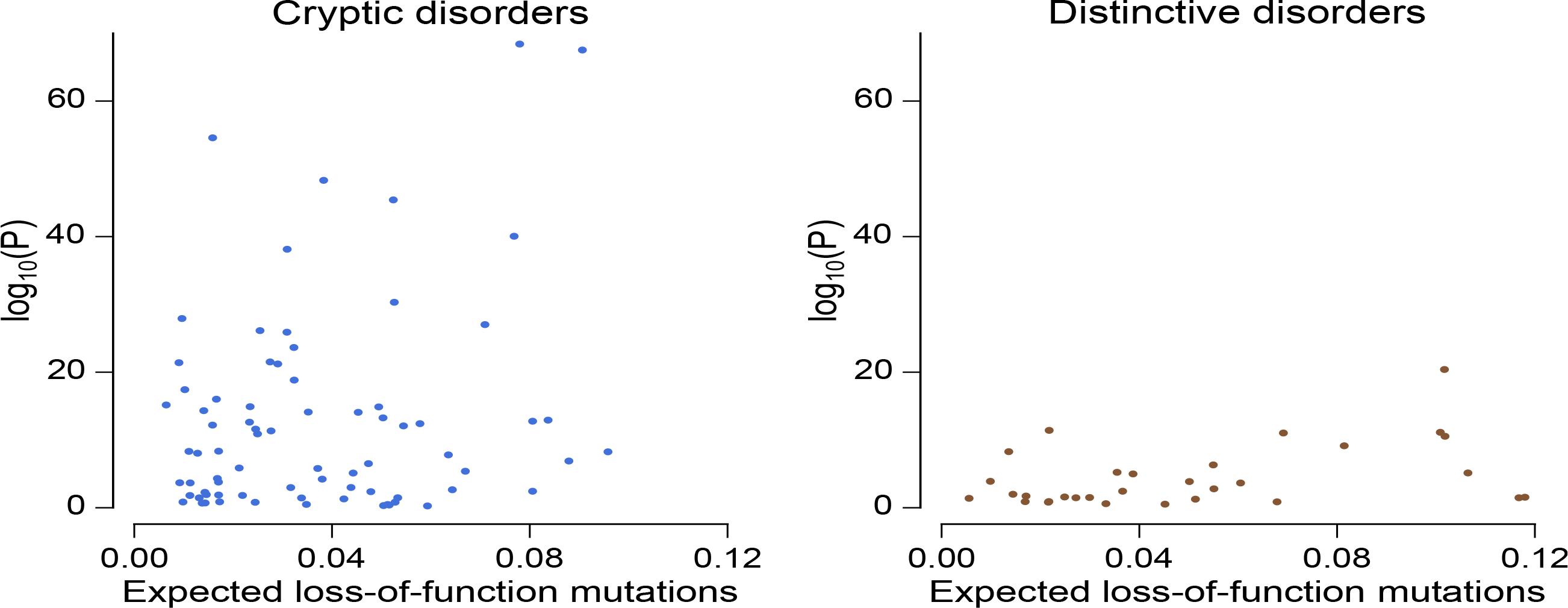
Neurodevelopmental genes classified by clinical recognisability were compared for the gene-wise significance versus the expected number of mutations per gene. Points are shaded by recognisability category. Genes have been separated into two plots, one plot with genes for cryptic disorders with low, mild or moderate clinical recognisability, and one plot with genes for distinctive disorders with high clinical recognisability.

**Supplementary Figure 8:**
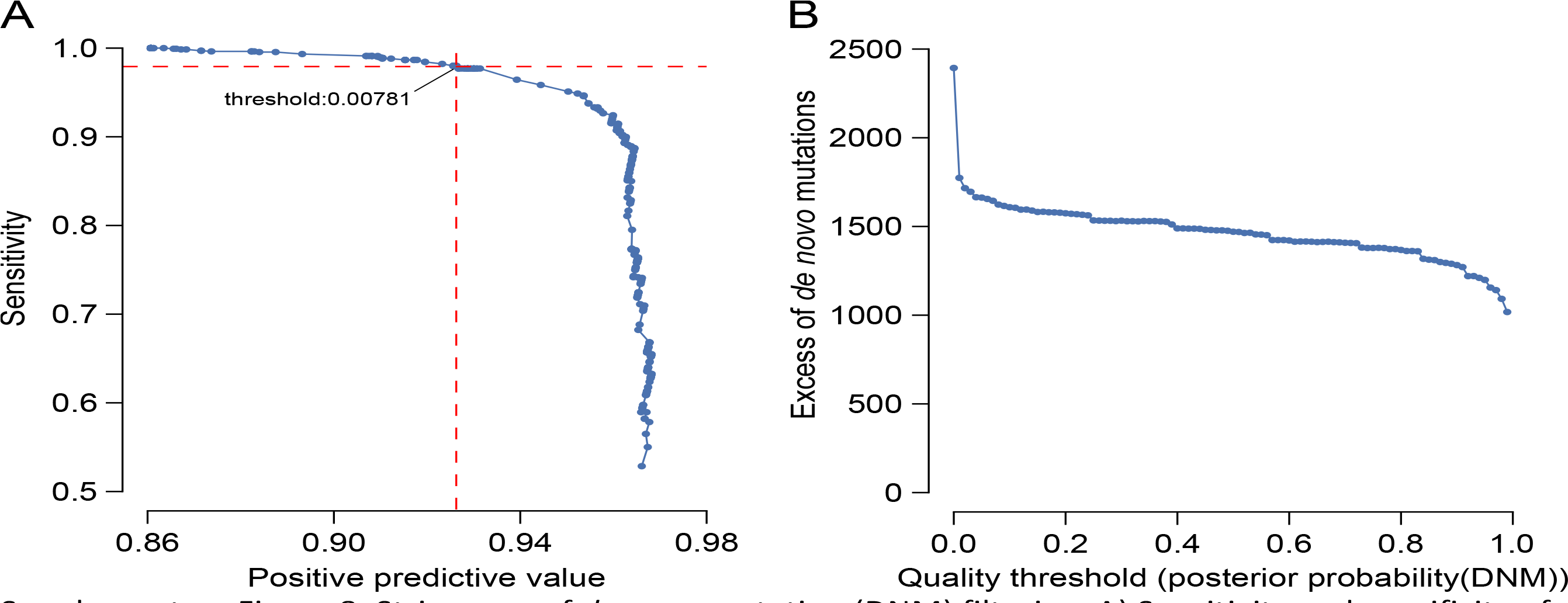
Stringency of *de novo* mutation (DNM) filtering. A) Sensitivity and specificity of DNM validations within sets filtered on varying thresholds of DNM quality (posterior probability of DNM). The analysed DNMs were restricted to sites identified within the earlier 1133 trios^15^, where all candidate DNMs underwent validation experiments. The labelled value is the quality threshold at which the number of candidate synonymous DNMs equals the number of expected synonymous mutations under a null germline mutation rate. B) Excess of missense and loss-of-function DNMs at varying DNM quality thresholds. The DNM excess is adjusted for the sensitivity and specificity at each threshold.

## References

1. Sheridan,E. et al. Risk factors for congenital anomaly in a multiethnic birth cohort: an analysis of the Born in Bradford study. Lancet 382, 1350–9 (2013).

2. Ropers,H.H. Genetics of early onset cognitive impairment. Annu Rev Genomics Hum Genet 11, 161–87(2010).

3. De Ligt,J. et al. Diagnostic exome sequencing in persons with severe intellectual disability. The New England Journal of Medicine 367, 1921–9 (2012).

4. De Rubeis,S. et al. Synaptic, transcriptional and chromatin genes disrupted in autism. Nature 515, 209–215 (2014).

5. Epi4K Consortium & Epilepsy Phenome/Genome Project. De novo mutations in epileptic encephalopathies. Nature 501, 217–21 (2013).

6. EuroEPINOMICS-RES Consortium, Epilepsy Phenome/Genome Project & Epi4K Consortium. De novo mutations in synaptic transmission genes including DNM1 cause epileptic encephalopathies. Am J Hum Genet 95, 360–70 (2014).

7. Fromer,M. et al. De novo mutations in schizophrenia implicate synaptic networks. Nature 506, 179–184 (2014).

8. Gilissen,C. et al. Genome sequencing identifies major causes of severe intellectual disability. Nature 511, 344–7 (2014).

9. Iossifov,I. et al. The contribution of de novo coding mutations to autism spectrum disorder. Nature 515, 216–221 (2014).

10. Iossifov,I. et al. De Novo Gene Disruptions in Children on the Autistic Spectrum. Neuron 74, 285–299 (2012).

11. O'Roak,B.J. et al. Sporadic autism exomes reveal a highly interconnected protein network of de novo mutations. Nature 485, 1–7 (2012).

12. Rauch,A. et al. Range of genetic mutations associated with severe non-syndromic sporadic intellectual disability: an exome sequencing study. Lancet 380, 1674–82 (2012).

13. Sanders,S.J. et al. De novo mutations revealed by whole-exome sequencing are strongly associated with autism. Nature 485, 237–41 (2012).

14. Zaidi,S. et al. De novo mutations in histone-modifying genes in congenital heart disease. Nature 498, 220–3 (2013).

15. The Deciphering Developmental Disorders Study. Large-scale discovery of novel genetic causes of developmental disorders. Nature 519, 223–228 (2015).

16. Wilkie,A.O. The molecular basis of genetic dominance. J Med Genet 31, 89–98 (1994).

17. Wright,C.F. et al. Genetic diagnosis of developmental disorders in the DDD study: a scalable analysis of genome-wide research data. The Lancet (2014).

18. Jacquemont,S. et al. A higher mutational burden in females supports a "female protective model" in neurodevelopmental disorders. Am J Hum Genet 94, 415–25 (2014).

19. Kong,A. et al. Rate of de novo mutations and the importance of father's age to disease risk. Nature 488, 471–5 (2012).

20. Rahbari,R. et al. Timing, rates and spectra of human germline mutation. Nat Genet 48, 126–33 (2016).

21. Wong,W.S. et al. New observations on maternal age effect on germline de novo mutations. Nat Commun 7, 10486 (2016).

22. Samocha,K.E. et al. A framework for the interpretation of de novo variation in human disease. Nature Genetics 46, 944–950 (2014).

23. Hirata,H. et al. ZC4H2 mutations are associated with arthrogryposis multiplex congenita and intellectual disability through impairment of central and peripheral synaptic plasticity. Am J Hum Genet 92, 681–95 (2013).

24. Homan,C.C. et al. Mutations in USP9X are associated with X-linked intellectual disability and disrupt neuronal cell migration and growth. Am J Hum Genet 94, 470–8 (2014).

25. Liu,J. et al. SMC1A expression and mechanism of pathogenicity in probands with X-Linked Cornelia de Lange syndrome. Hum Mutat 30, 1535–42 (2009).

26. Akawi,N. et al. Discovery of four recessive developmental disorders using probabilistic genotype and phenotype matching among 4,125 families. Nature Genetics 47, 1363–1369 (2015).

27. Meynert,A.M., Ansari,M., FitzPatrick,D.R. & Taylor,M.S. Variant detection sensitivity and biases in whole genome and exome sequencing. BMC Bioinformatics 15, 247 (2014).

28. Lek,M. et al. Analysis of protein-coding genetic variation in 60,706 humans. bioRxiv X, XX–XX (2015).

29. Boycott,K.M., Vanstone,M.R., Bulman,D.E. & Mackenzie,A.E. Rare-disease genetics in the era of next-generation sequencing: discovery to translation. Nature Reviews Genetics 14, 681–91 (2013).

30. Springett,A. et al. Congenital Anomaly Statistics 2011: England and Wales. (2013).

31. Bragin,E. et al. DECIPHER: database for the interpretation of phenotype-linked plausibly pathogenic sequence and copy-number variation. Nucleic Acids Res 42, D993–D1000 (2014).

32. Kohler,S. et al. Clinical diagnostics in human genetics with semantic similarity searches in ontologies. American Journal of Human Genetics 85, 457–464 (2009).

33. Li,H. & Durbin,R. Fast and accurate short read alignment with Burrows-Wheeler transform. Bioinformatics 25, 1754–1760 (2009).

34. McKenna,A. et al. The Genome Analysis Toolkit: a MapReduce framework for analyzing next-generation DNA sequencing data. Genome Res 20, 1297–303 (2010).

35. Li,H. et al. The Sequence Alignment/Map format and SAMtools. Bioinformatics 25, 2078–2079 (2009).

36. Ramu,A. et al. DeNovoGear: de novo indel and point mutation discovery and phasing. Nature Methods 10, 985–7 (2013).

37. Abecasis,G.R. et al. An integrated map of genetic variation from 1,092 human genomes. Nature 491, 56–65 (2012).

38. McLaren,W. et al. Deriving the consequences of genomic variants with the Ensembl API and SNP Effect Predictor. Bioinformatics 26, 2069–70 (2010).

39. Felzenszwalb,P.F., Girshick,R.B., McAllester,D. & Ramanan,D. Object detection with discriminatively trained part-based models. IEEE transactions on pattern analysis and machine intelligence 32, 1627–45 (2010).

40. Xiong,X. & De la Torre,F. Supervised Descent Method and Its Applications to Face Alignment. in 2013 IEEE Conference on Computer Vision and Pattern Recognition (CVPR) 532–539 (IEEE, Portland, OR, 2013).

41. Ferry,Q. et al. Diagnostically relevant facial gestalt information from ordinary photos. eLife 3, e02020–e02020 (2014).

42. Cooper,G.M. et al. A copy number variation morbidity map of developmental delay. Nat Genet 43, 838–46 (2011).

43. Sagoo,G.S. et al. Array CGH in patients with learning disability (mental retardation) and congenital anomalies: updated systematic review and meta-analysis of 19 studies and 13,926 subjects. Genet Med 11, 139–46 (2009).

